# A Lineage-Specific Peptide Suppresses Juvenile Hormone to Drive Reproductive and Longevity Reprogramming in Ants

**DOI:** 10.64898/2026.05.23.727378

**Authors:** Long Ding, Jakub Mlejnek, Haiyan Zheng, Ching-Han Lee, Yen-Chung Chen, Danny Reinberg, Claude Desplan

**Affiliations:** Department of Biology, New York University; New York, NY 10003, USA; Biological Mass Spectrometry Facility, Robert Wood Johnson Medical School and Rutgers, The State University of New Jersey; Piscataway, NJ 08854, USA; Department of Human Genetics, Miller School of Medicine, University of Miami; Miami, FL 33136, USA.; Howard Hughes Medical Institute, Miller School of Medicine, University of Miami; Miami, FL 33136, USA

## Abstract

In ants and other eusocial insects, reproductive division of labor is tightly regulated by juvenile hormone (JH), which suppresses reproduction in most individuals to maintain the worker caste. However, how social cues trigger systemic JH suppression to permit reproductive activation remains unclear. In the ant *Harpegnathos saltator*, workers respond to queen loss by engaging in sustained ritualistic dueling and transitioning into reproductive, long-lived pseudoqueens (gamergates). This provides a powerful model for investigating the molecular basis of socially induced plasticity. We examined the hemolymph proteome of transitioning ants and identified HCRG1, a lineage-restricted peptide upregulated during dueling, as a circulating factor that physically interacts with Hex70c, a broadly conserved JH-binding Hexamerin. HCRG1 promotes ovarian development by antagonizing JH signaling. Expression of ant *HCRG1* in the heterologous solitary model *Drosophila* also extends lifespan. These findings identify an evolutionarily derived circulating factor that links social sensory perception to systemic JH suppression, enabling coordinated reproductive and longevity transitions in a social insect.

## INTRODUCTION

Unlike most eusocial insects, the Indian jumping ant *Harpegnathos saltator* retains remarkable developmental plasticity into adulthood.^1–3^. In the absence of queen pheromones, genetically similar, non-reproductive female workers engage in dueling behavior^2,4^. Although most workers rapidly abandon the duel, a small number of winners continue dueling for more than 30 days to become stable reproductive individuals, known as gamergates, which maintain the colony. Early sustained dueling activity serves as a strong predictor of an eventual transformation into gamergates, with ovarian development observable in persistent duelers by day 3 after initiation of dueling, and egg-laying beginning between days 6 and 10^2^. Alongside behavioral and reproductive changes, gamergates exhibit significant metabolic alterations for oogenesis and a dramatic fivefold increase in lifespan^5,6^.

In established colonies, dueling and the worker-to-gamergate caste transition are repressed by queen pheromones, long-chain cuticular hydrocarbons, that activate specific olfactory receptors located on antennal sensilla^7–9^. Disruption of olfactory input, such as by mutation of the olfactory co-receptor *orco*, prevents dueling and blocks gamergate formation^1^. Exposure to queen pheromones leads to an increase in the brain of neuropeptide Corazonin (Crz), which promotes worker behaviors such as foraging, and maintains workers in a non-reproductive state^10^. The transition to the gamergate state is accompanied by profound physiological changes, including brain shrinkage and alterations in neural composition and activity^11–13^, along with major endocrine shifts: the levels of JH III decline, while 20-hydroxyecdysone (20E) increases^14,15^. JH has emerged as a central player in eusocial evolution^16,17^: the fine-tuned regulation of JH is required for labor division in ant societies. Although JH promotes reproduction in most insect species, it suppresses reproduction in worker ants. As a consequence, the suppression of JH is essential to allow reproductive development in queens or gamergates^14,18^. For example, exogenous JH III suppresses ovary development and reinforces worker-like behavior, whereas 20E promotes reproductive traits^2,14^. However, the mechanism by which olfactory input in the brain is transduced into systemic JH suppression remains unresolved.

To address this, we profiled the hemolymph proteome of non-dueling and dueling workers and identified HCRG1, a small Kunitz-domain peptide that is upregulated during early caste transition. This peptide, previously found to be transcriptionally enriched in the brain of early duelers and gamergates^2^, is required for both behavioral and reproductive aspects of gamergate formation. Using immunoprecipitation followed by mass spectrometry (IP-MS), we found that HCRG1 binds to the hemolymph hexameric protein Hex70c, a member of the Hexamerin family known to bind directly to JH^16,19^. Comparative evolutionary analyses further indicate that HCRG1 is a lineage-restricted and rapidly evolving gene in ant-associated lineages, whereas Hex70c is broadly conserved across insects. Structural modeling suggests that this interaction remodels the ligand-binding pocket of Hex70c, enhancing its ability to sequester JH and thereby reduce JH signaling. In accordance, *HCRG1* knockdown (KD) in duelers disrupts the JH-responsive transcriptomic program and arrests ovarian maturation, while supplementation with recombinant HCRG1 overrides the inhibitory effects of exogenous JH and restores reproductive caste transition. Furthermore, HCRG1 overexpression in *Drosophila* extends female lifespan, mimicking the known effects of JH depletion^20–22^. Together, our findings reveal a previously unrecognized brain-to-body mechanism in which an evolutionarily derived circulating peptide engages a conserved Hexamerin to suppress JH signaling, thereby enabling reproductive caste transition and associated longevity plasticity in ants.

## RESULTS

### Changes in the hemolymph proteome during caste transition

Dueling behavior within the first 3 days of the tournament is an early indicator of caste fate, with high-level dueling predicting gamergate transition and low or no dueling associated with worker identity^2^. Our previous studies have identified a set of brain markers associated with antennal dueling and caste determination that emerge in the early stages of transition^2^. To determine whether such brain-derived factors are released into the circulation to regulate broad organismal reprogramming, we performed quantitative proteomic analysis of hemolymph, the open circulatory system of insects, which was collected from ants classified as duelers and non-duelers at four time points: 0, 3, 10, and 30 days after the onset of the tournament (four biological replicates per group). Proteomic analysis identified 649 proteins across all hemolymph samples. Comparative analysis between duelers and non-duelers at each time point revealed a progressive increase in the number of differentially expressed proteins (DEPs), suggesting that hemolymph composition diverges over time as individuals progress through the caste transition (**Fig. 1a**). To identify brain-derived factors that may act as systemic regulators during early transformation, we integrated these data with our previously published brain transcriptome. This analysis revealed four factors that were consistently upregulated in the brain and were also detected as DEPs in the hemolymph of day 10 duelers: the Kunitz-type serine protease inhibitor HCRG1 and three uncharacterized proteins. Structural prediction using the MPI Toolkit suggested that LOC105181259 was a putative Apolipophorin-III, LOC105189919 was an odorant binding protein, and LOC109504028 was a chitinase (**Fig. 1b and Extended Data Fig. 1a**). LOC105189919 was also significantly upregulated in the fat body of day 10 duelers (**Extended Data Fig. 1, b and d**), while HCRG1 and LOC109504028 were both transcriptionally downregulated in the ovary (**Extended Data Fig. 1c and e**).

**Fig. 1.**
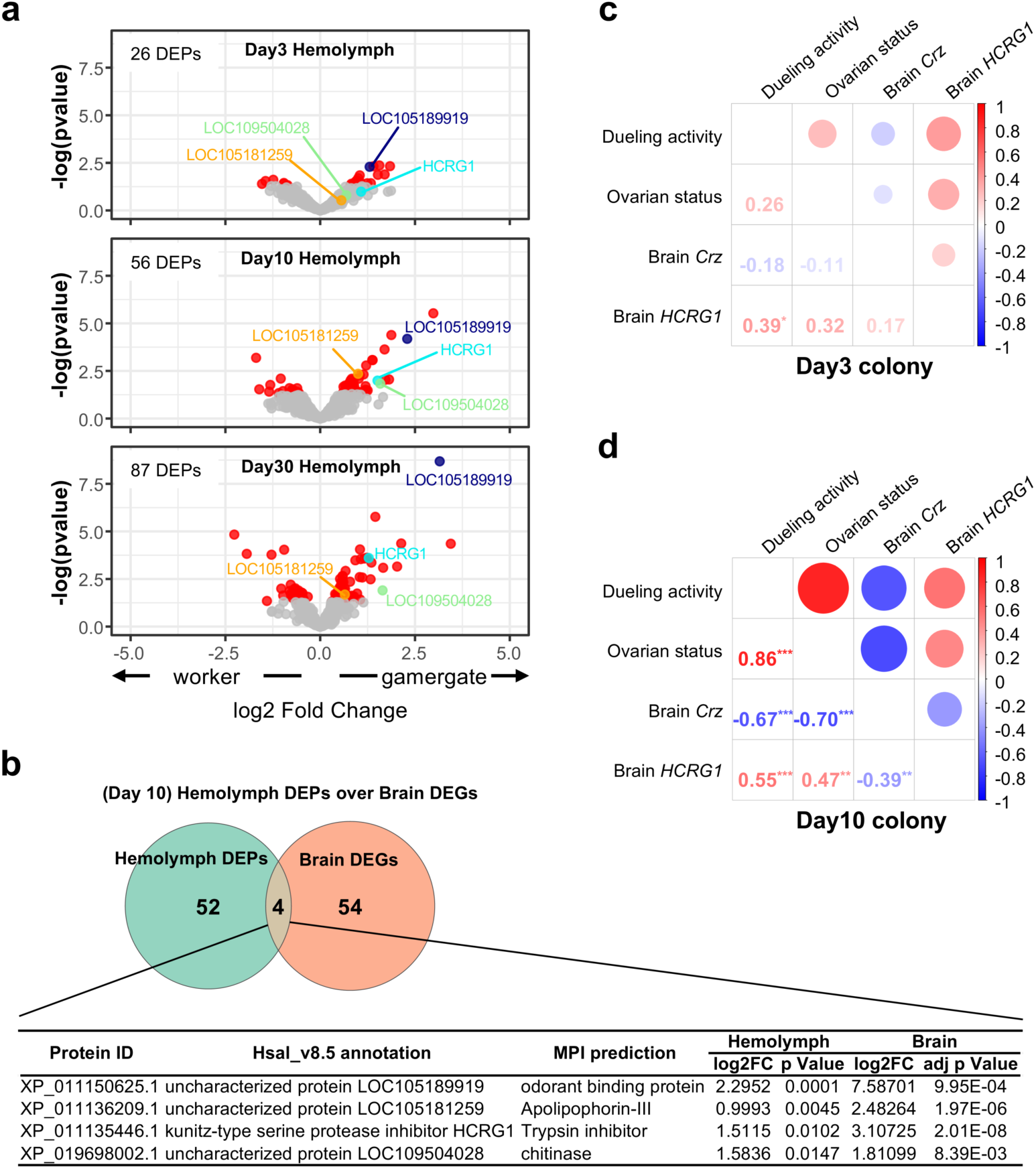
Divergence of hemolymph proteomes during caste transition. **a,** Volcano plot of Tandem Mass Tag (TMT) MS from hemolymph of duelers and non-duelers at different time points during early caste transition. Each circle represents a protein. Significant proteins (p < 0.05) are shown in red, while nonsignificant proteins (p > 0.05) are shown in gray. HCRG1 (cyan), LOC105181259 (orange), LOC105189919 (navy), and LOC109504028 (green) are highlighted. The number of DEPs identified at each time point is indicated in the upper left corner of each plot. **b,** Venn diagram comparing the central brain transcriptome and hemolymph proteome at day 10. The four shared factors are listed below the diagram. **c,d,** Correlation analysis of dueling activity, ovarian development, and brain gene expression in ants from day 3 (**c**) and day 10 (**d**) colonies. Circle size represents the magnitude of the absolute correlation coefficient. Numbers indicate the specific correlation coefficient values. Colors: Blue indicates negative correlation, red indicates positive correlation. Asterisks denote significance thresholds for each correlation: * for 0.01 < p ≤ 0.05, ** for 0.001 < p ≤ 0.01, and *** for p ≤ 0.001.

### HCRG1 is a marker of duelers

We had previously identified HCRG1 as a dueler-associated marker in the brain^2^. Its detection in the hemolymph proteome suggested a potential systemic role in caste differentiation. To investigate the association between brain *HCRG1* expression and caste transition, we analyzed all individuals exhibiting varying levels of dueling activity from a transitioning colony. *HCRG1* expression in the brain was positively correlated with both dueling frequency and the presence of yolky oocytes. This correlation was more pronounced in day 10 individuals compared to day 3, consistent with the notion that prolonged dueling promotes the acquisition of gamergate fate (**Fig. 1c and d**). In contrast, brain expression of *Crz*, a canonical marker of the worker state^10^, was negatively correlated with dueler-associated traits, exhibiting an inverse trend relative to *HCRG1* (**Fig. 1c and d**).

### HCRG1 is required for gamergate transition

To investigate the role of HCRG1 during the worker-to-gamergate transition, we performed loss-of-function experiments by injecting *HCRG1* dsRNA into the heads of day 3 duelers and tracking their phenotypic changes over 4 days (**Fig. 2a**). *HCRG1* mRNA expression in the brain was significantly reduced in a dose-dependent manner (**Extended Data Fig. 2a**). Although transcriptomic data suggest that the fat body also produces HCRG1, its expression remained largely unaffected by head injections (**Extended Data Fig. 2b**). Consistent with its reduced expression in the brain, hemolymph HCRG1 protein levels declined in a dose-dependent manner following injection (**Fig. 2b and c**), suggesting that its primary source is the brain of duelers.

**Fig. 2.**
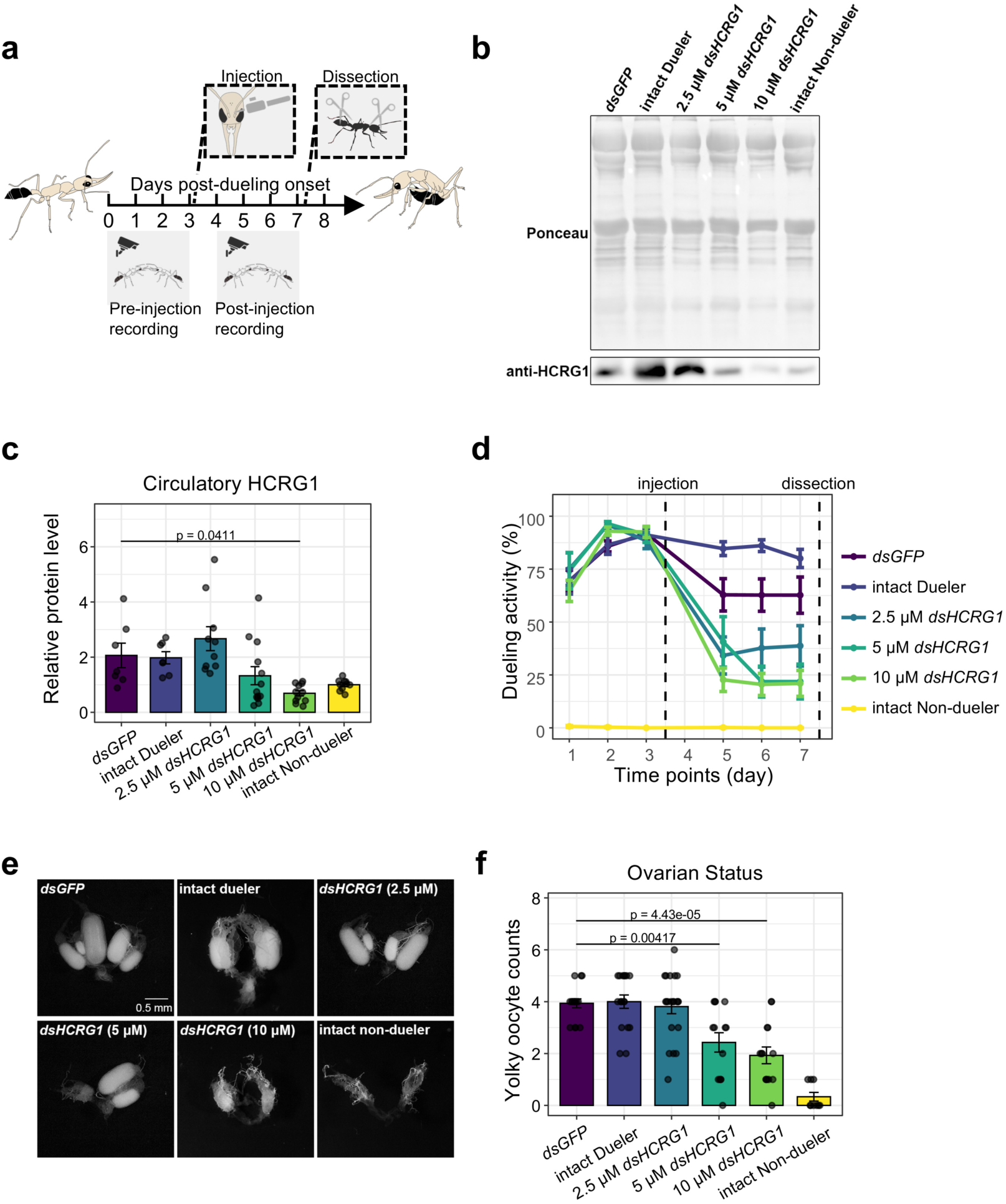
HCRG1 knockdown inhibits worker-to-gamergate transition. **a,** General schematic of the injection-based treatment paradigm used in this study. Individuals at day 3 of the caste transition were subjected to injection, and samples were collected on the fourth day post-injection for downstream analysis. Each tick mark along the timeline arrow represents one day. **b,** Western blot analysis of representative hemolymph samples from different treatments. Ponceau staining of the entire membrane (top) and anti-HCRG1 bands (bottom) are shown. **c,** Quantification of relative circulatory HCRG1 protein levels. The band intensity of anti-HCRG1 was normalized to the Ponceau intensity of the corresponding lane. Sample sizes: *dsGFP* (n = 7), intact duelers (n = 7), 2.5 μM *dsHCRG1* (n = 10), 5 μM *dsHCRG1* (n = 13), 10 μM *dsHCRG1* (n = 11), and intact non-duelers (n = 10). Results from multiple colonies were combined and each ratio was normalized to the average value of the non-duelers from the same colony. Error bars represent SEM. *p*-value from one-way ANOVA with Tukey’s post hoc analysis. **d,** Quantification of dueling activity. Error bars represent SEM. Sample sizes: *dsGFP* (n = 6), intact duelers (n = 7), 2.5 μM *dsHCRG1* (n = 20), 5 μM *dsHCRG1* (n = 16), 10 μM *dsHCRG1* (n = 15), and intact non-duelers (n = 11). **e,** Representative images of dissected ovaries 4 days post-injection (scale bar = 0.5 mm). **f,** Quantification of yolky oocytes. Sample sizes: *dsGFP* (n = 16), intact duelers (n = 17), 2.5 μM *dsHCRG1* (n = 21), 5 μM *dsHCRG1* (n = 14), 10 μM *dsHCRG1* (n = 14), and intact non-duelers (n = 9). Error bars represent SEM. *p*-values from one-way ANOVA with Tukey’s post hoc analysis.

Behavioral assays revealed that while *GFP* RNAi control ants showed a mild reduction in dueling activity likely due to the stress of injection in the head (**Fig. 2d**), individuals treated with *HCRG1* RNAi and exhibiting low circulating HCRG1 levels displayed a significantly stronger suppression of dueling behavior (**Fig. 2d and Extended Data Fig. 2c**). Correspondingly, ovarian development at day 7 was arrested at the level of day 3-dueler stage (**Fig. 2, e-f and Extended Data Fig. 2e-f**) and failed to progress toward gamergate physiology. Thus, we observed strong positive correlations between brain *HCRG1* expression, hemolymph HCRG1 levels, ovarian development, and dueling activity (**Extended Data Fig. 2c**). While brain *Crz* expression was upregulated in all injected ants compared to intact duelers, it did not differ significantly between the *GFP* RNAi and *HCRG1* RNAi groups (**Extended Data Fig. 2d**), suggesting a general stress response rather than a specific effect of *HCRG1* KD. Together, these findings suggest that brain-derived circulatory HCRG1 regulates caste transition and is essential for sustaining dueling behavior and for reproductive activation in gamergates.

### Elevation of circulating HCRG1 induces ovarian activation

To determine whether the phenotypic effects of *HCRG1* RNAi resulted from reduced circulating HCRG1, we performed rescue experiments using recombinant HCRG1 protein expressed in Sf9 cells. Mass spectrometry analysis identified histidine19 as the start site of the secreted form of HCRG1 after the processing of its signal peptide. To facilitate recombinant expression and purification, we inserted a 6×His tag followed by a TEV protease recognition site between the signal peptide and the secreted peptide (**Fig. 3a**). The purified protein with its His-Tag was used to generate polyclonal antibodies, while the circulating recombinant protein fragment (RP), which mimics the endogenous secreted form, was isolated after TEV digestion for use in rescue experiments (**Fig. 3a**).

**Fig. 3.**
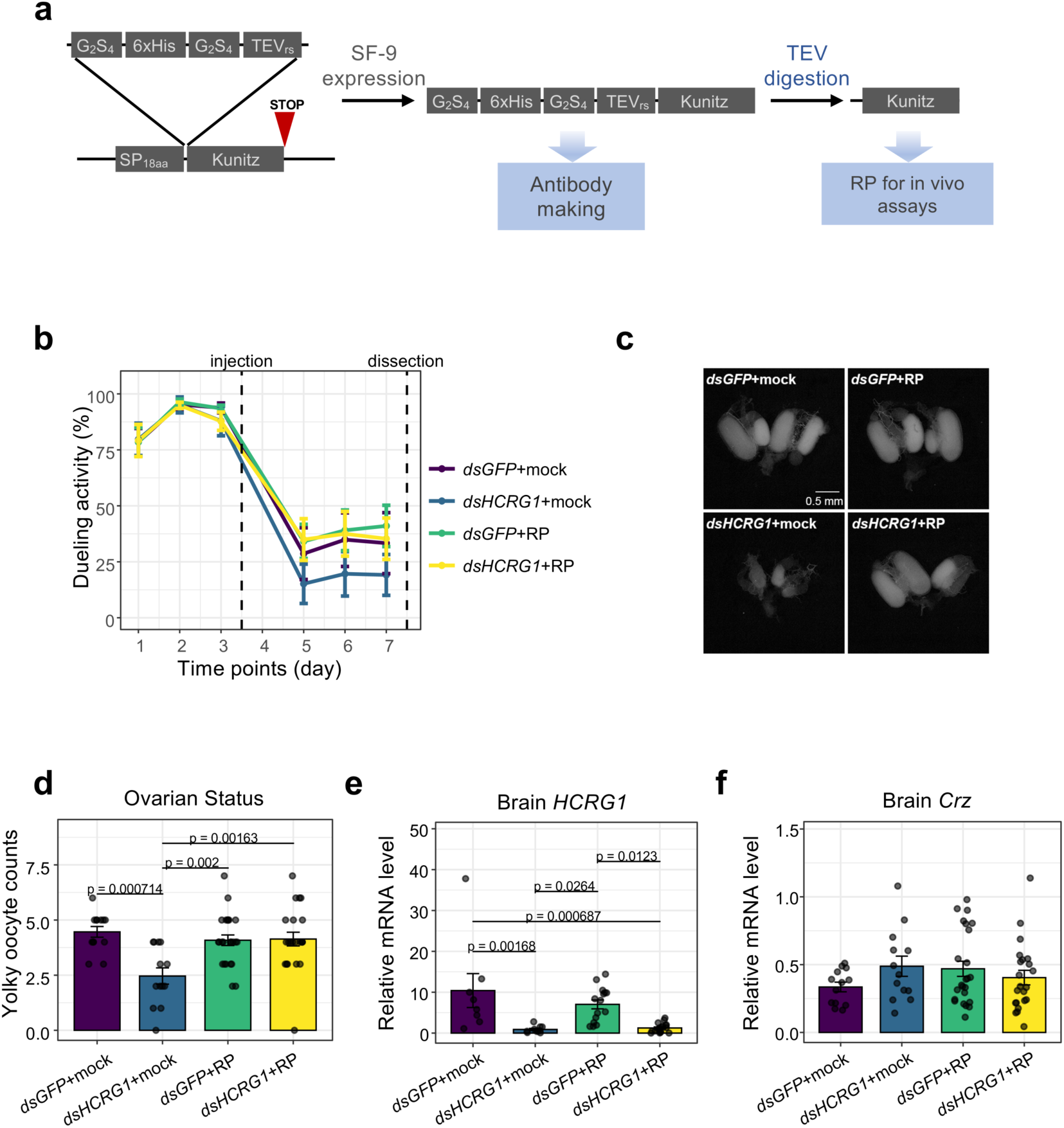
Rescue of HCRG1 knockdown defects by recombinant protein. **a,** Schematic of the HCRG1 recombinant protein design. A 6×His tag flanked by G_2_S_4_ linkers and a TEV recognition site were inserted into the HCRG1 protein, downstream of the 18 amino acid N-terminal signal peptide (SP). Purified protein with these elements was used for antibody production. After TEV cleavage, the Kunitz domain-containing fragment was used for recombinant protein administration assays. **b,** Quantification of dueling activity. Error bars represent SEM. *dsGFP*+mock (n = 13), *dsHCRG1*+mock (n = 13), *dsGFP*+RP (n = 24), *dsHCRG1*+RP (n = 22). **c,** Representative images of dissected ovaries 4 days post-injection (scale bar = 0.5 mm). **d,** Quantification of yolky oocytes. Error bars represent SEM. *p*-values from one-way ANOVA with Tukey’s post-hoc analysis. *dsGFP*+mock (n = 13), *dsHCRG1*+mock (n = 13), *dsGFP*+RP (n = 24), *dsHCRG1*+RP (n = 22). **e-f,** qPCR analysis of *HCRG1* (**e**) and *Crz* (**f**) expression in brains from different treatment groups. *RPL32* was used as an internal control. Results from multiple colonies were combined, and each expression value was normalized to the average of non-duelers from the same colony. Error bars indicate SEM. Statistical significance was determined by one-way ANOVA followed by Tukey’s post hoc analysis. Sample sizes: **e,** *dsGFP*+mock (n = 8), *dsHCRG1*+mock (n = 15), *dsGFP*+RP (n = 10), *dsHCRG1*+RP (n = 18); **f,** *dsGFP*+mock (n = 13), *dsHCRG1*+mock (n = 13), *dsGFP*+RP (n = 24), *dsHCRG1*+RP (n = 22).

To restore circulating HCRG1 levels, we injected RP into the abdomen of ants treated with *HCRG1* RNAi in the head, ensuring direct hemolymph delivery. This treatment rescued both dueling activity and ovarian development in *HCRG1* RNAi duelers, restoring them to control levels (**Fig. 3, b-d**). Furthermore, although RP injection in day 3 non-duelers induced ovarian development after 4 days, it did not trigger dueling behavior (**Extended Data Fig. 3, c and d**). Moreover, the expression of *HCRG1* in the brain of rescued RNAi duelers remained suppressed (**Fig. 3e**), confirming that RP supplementation, rather than insufficient knockdown efficiency, led to the rescue effect. Additionally, brain *Crz* expression was unchanged between RP-rescued and non-rescued duelers (**Fig. 3f**), indicating that Crz is unlikely to function downstream of HCRG1. In RP-injected non-duelers in which ovarian development was induced, neither brain *HCRG1* nor *Crz* expression were significantly altered (**Extended Data Fig. 3a and b**), further supporting the notion that circulating HCRG1 directly promotes ovarian activation without modifying brain *Crz* levels. These findings show that circulatory HCRG1 is both necessary and sufficient to activate reproduction. It is also necessary, but not sufficient for the induction of dueling.

### HCRG1 interacts with a hexamerin protein

To investigate the molecular mechanisms through which HCRG1 functions, we performed immunoprecipitation followed by mass spectrometry (IP-MS) on whole-ant lysates from day 10 duelers and non-duelers, using an HCRG1-specific antibody to identify its interacting partners during caste transition. This analysis identified a highly abundant hemolymph protein annotated as arylphorin subunit alpha isoform X1 (**Extended Data Fig. 1a**), encoded by the *hexamerin 70c* (*Hex70c*) gene. This interaction was detected independently in both duelers and non-duelers (**Fig. 4a and Extended Data Fig. 4a**). Hexamerin has been shown to function as a JH-binding protein in insects^23^. In paper wasps, hexamerins silence JH signaling to promote gyne potentiation^24^, while in termites, direct binding to JH sequesters the hormone and limits worker-to-soldier transition^19,25,26^. Thus, HCRG1 may regulate JH signaling through its interaction with Hex70c. Moreover, the upregulation of HCRG1 in duelers correlates with a reduction in JH titers^14,15^, further supporting the hypothesis that the HCRG1-Hexamerin interaction is involved in the suppression of JH activity during caste transition.

**Fig. 4.**
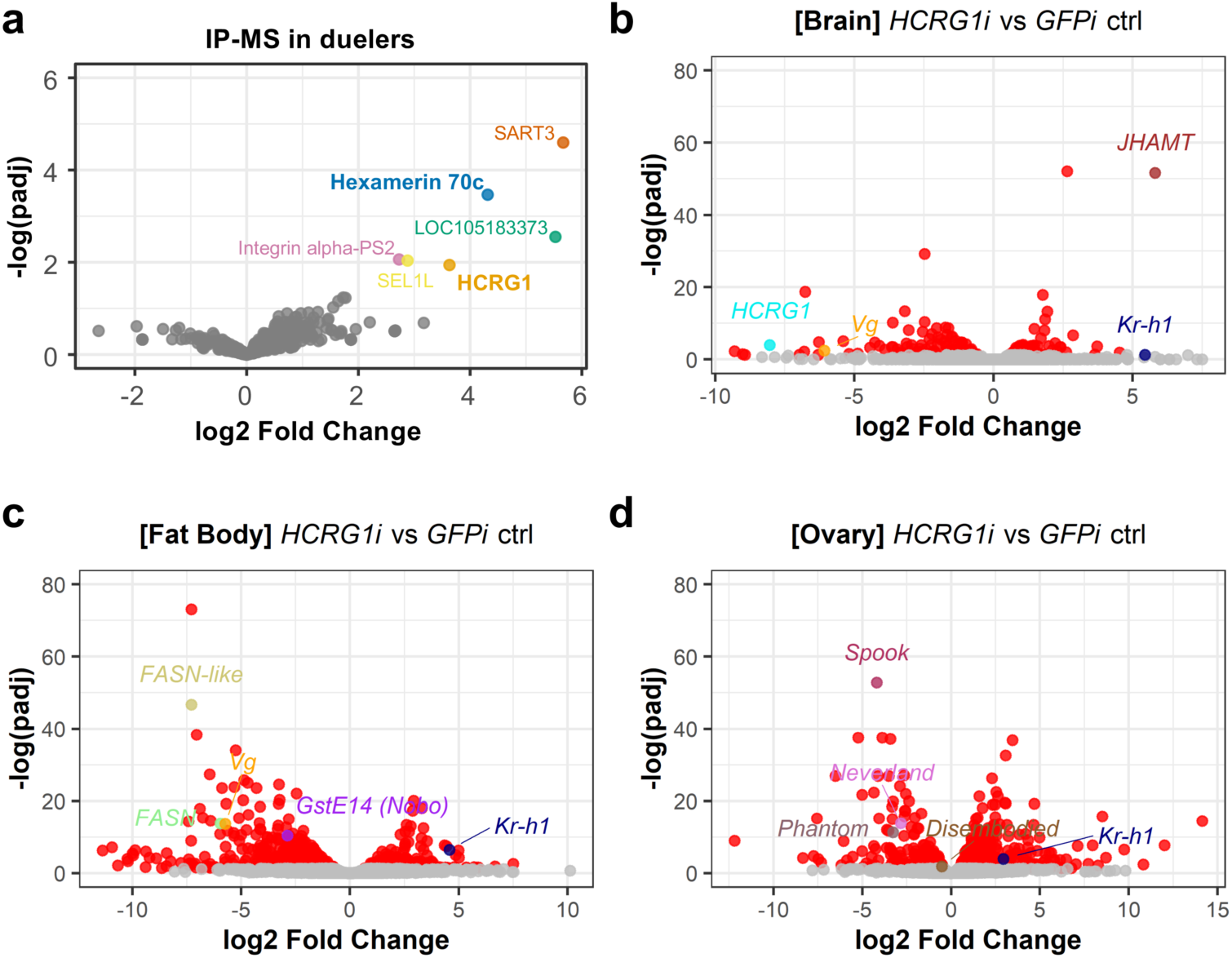
Proteins interacting with HCRG1 and its downstream targets. **a,** Volcano Plot of IP-MS from duelers. Proteins enriched in HCRG1 antibody-crosslinked beads were compared to the IgG control. Each dot represents an identified protein. Proteins significantly enriched with the HCRG1 antibody (adjusted p < 0.05) are highlighted in colors: Hexamerin 70c (blue), HCRG1 (orange), SART3 (vermilion), LOC105183373 (green), Integrin alpha-PS2 (magenta), and SEL1L (yellow). Non-specific binding proteins (adjusted p > 0.05) are shown in gray. **b-d,** Volcano Plots of bulk RNA-Seq from brain (**b**), fat body (**c**), and ovary (**d**). Transcriptomes from *HCRG1* RNAi duelers (*HCRG1i*) were compared to *GFP* RNAi controls (*GFPi* ctrl). Differentially expressed genes (DEGs, adjusted p < 0.05) are shown in red, while non-significant genes (adjusted p > 0.05) are shown in gray. Selected genes of interest are color-coded and labeled: *JHAMT* (brown), *HCRG1* (cyan), *Vg* (orange), *Kr-h1* (navy), *FASN-like* (khaki), *FASN* (light green), *GstE14 (Nobo)* (purple), *Spook* (maroon), *Neverland* (orchid), *Phantom* (dark pink), and *Disembodied* (tan).

To further evaluate this interaction, we employed AlphaFold3 (AF3) structural modeling that yielded high-confidence predictions (ipTM = 0.92, pTM = 0.93) for the HCRG1-Hex70c complex (**Extended Data Fig. 6a**), supporting the likelihood of a direct protein-protein interaction *in vivo*. We next asked whether HCRG1 binding influences the capacity of Hex70c to associate with JH III. Using AutoDock Vina^27^, we first benchmarked the docking workflow against the experimentally resolved silkworm JH-binding protein–JH III complex^28^, which produced a predicted binding energy of −8.435 ± 0.013 kcal/mol (**Extended Data Fig. 7a**). Iterative global-to-local docking of JH III to the predicted Hex70c hexamer identified a best predicted score of approximately −8.519 kcal/mol in the absence of HCRG1. Although docking was performed with a single JH III molecule, the HCRG1–Hex70c complex contained six geometrically symmetric internal pockets, suggesting that the hexameric assembly could accommodate JH III in multiple equivalent sites (**Extended Data Fig. 6a**). To test whether the improved score reflected the HCRG1-bound conformation rather than pocket position alone, we remapped the best HCRG1-bound JH III pose onto the unbound Hex70c structure and re-docked it. This reduced the predicted binding score to −8.485 kcal/mol (**Extended Data Fig. 7a**), suggesting that HCRG1 allosterically enhances Hex70c’s binding capacity for JH III. Consistent with this interpretation, structural alignment of the HCRG1-bound and unbound Hex70c states revealed a substantial displacement of the docked JH III pose (RMSD ≈ 8.76 Å), whereas local rearrangements within the surrounding pocket were comparatively subtle (**Extended Data Fig. 7b**). Moreover, HCRG1 did not directly contact JH III in the predicted complex (**Extended Data Fig. 6b**), indicating that its effect is unlikely to result from direct ligand coordination. Together, these analyses suggest that HCRG1 may indirectly enhance the JH III-binding capacity of Hex70c, potentially promoting Hex70c-mediated sequestration or suppression of circulating JH through an allosteric mechanism.

### HCRG1 is implicated in the regulation of the JH pathway

The identification of the HCRG1-Hex70c interaction directed our attention to the JH pathway. To investigate how HCRG1 might influence the JH pathway, we performed bulk RNA-sequencing on brains, fat bodies, and ovaries from intact duelers and non-duelers as well as duelers subjected to head-injection RNAi treatments targeting either *GFP* (control) or *HCRG1* (**Fig. 2a**). Transcriptomic distance (TSD) analysis demonstrated that the *GFP* RNAi control group clustered closely with duelers but was transcriptionally distant from non-duelers across all three tissues (**Extended Data Fig. 4b–d**), validating its use as a control for filtering out injection-induced background effects. The degree of transcriptional disruption following *HCRG1* KD varied across tissues, with the most pronounced changes observed in the ovary, followed by the fat body, while the brain exhibited the least alteration (**Fig. 4b-d and Extended Data Fig. 4b-d**). This pattern aligned with the results of Gene Set Enrichment Analysis (GSEA) (**Extended Data Fig. 5**), suggesting that HCRG1 acts as a systemic regulator in peripheral tissues.

In the brain, RNAi delivered via head injection effectively reduced *HCRG1* expression in duelers (**Fig. 4b**). Notably, the most prominently upregulated gene was *JHAMT*, the enzyme that catalyzes the key step in JH biosynthesis and is subject to positive feedback from JH signaling^29,30^, further implicating HCRG1 in the regulation of this pathway. Consistent with this, *Krüppel homolog 1* (*Kr-h1*), a well-established JH responsive transcription factor^14,31^, was also upregulated upon *HCRG1* RNAi, whereas *Vitellogenin* (*Vg*), which is negatively regulated by JH in social insects^15,32,33^, was significantly downregulated (**Fig. 4b**). This suggests that *HCRG1* KD activates the JH pathway, which is normally suppressed in duelers with high circulating HCRG1.

In the fat body, *HCRG1* KD affected genes associated with lipid metabolism and fatty acid biosynthesis, an effect that was also observed in the ovary (**Extended Data Fig. 5**): multiple fatty acid synthase genes, such as *FASN-like*, *FASN*, and *Vg*, a key gene promoting vitellogenesis in the fat body, were significantly downregulated. *Vg* and these lipogenesis-related genes, including FAS, are known to be negatively regulated by JH^15,29,30,32–36^. As in the brain, *Kr-h1* was strongly upregulated, further reinforcing the model that systemic activation of JH signaling occurs in response to HCRG1 depletion. (**Fig. 4c**).

In the ovary, a global downregulation of ribosome biogenesis-related genes was observed (**Extended Data Fig. 5**), suggesting that *HCRG1* KD induced ovarian quiescence. The most significantly downregulated gene was *Spook* (*Spo*), a member of the Halloween gene family encoding key enzymes for 20E biosynthesis^37,38^. Additionally, other ovary-resident Halloween genes, including *Neverland* (*Nvd*), *Phantom* (*Phtm*), and *Disembodied*, were also downregulated following *HCRG1* RNAi (**Fig. 4d**). This is consistent with prior findings that Halloween gene expression can be directly suppressed by JH signaling through its downstream effector Kr-h1^31,39^. Notably, *GstE14*/*noppera-bo* (*Nobo*), another Halloween enzyme responsible for cholesterol intake, a crucial upstream step in 20E biosynthesis, was selectively downregulated in *HCRG1* RNAi fat bodies (**Fig. 4c**). Given that 20E biosynthesis is essential for female reproductive development^37,38,40,41^, the suppression of Halloween gene expression provides a mechanistic explanation for the defects in ovarian development observed upon *HCRG1* KD. These findings further reflect the known antagonistic relationship between JH and 20E for caste differentiation in *Harpegnathos*^14,15^.

These transcriptional responses across multiple tissues suggest that HCRG1 functions as a systemic inhibitor of JH signaling, potentially through interaction with Hex70c.

### HCRG1 counteracts JH to drive social gamergate formation

To explore whether the upregulation of HCRG1 in early duelers functions by antagonizing JH signaling, we investigated its interaction with JH. Head injection of JH III into day 3 duelers significantly suppressed both dueling intensity and ovarian maturation (**Fig. 5a-c**). However, these inhibitory effects were reversed when HCRG1 RP was co-administered, indicating that HCRG1 prevents JH III from suppressing gamergate phenotypes.

**Fig. 5.**
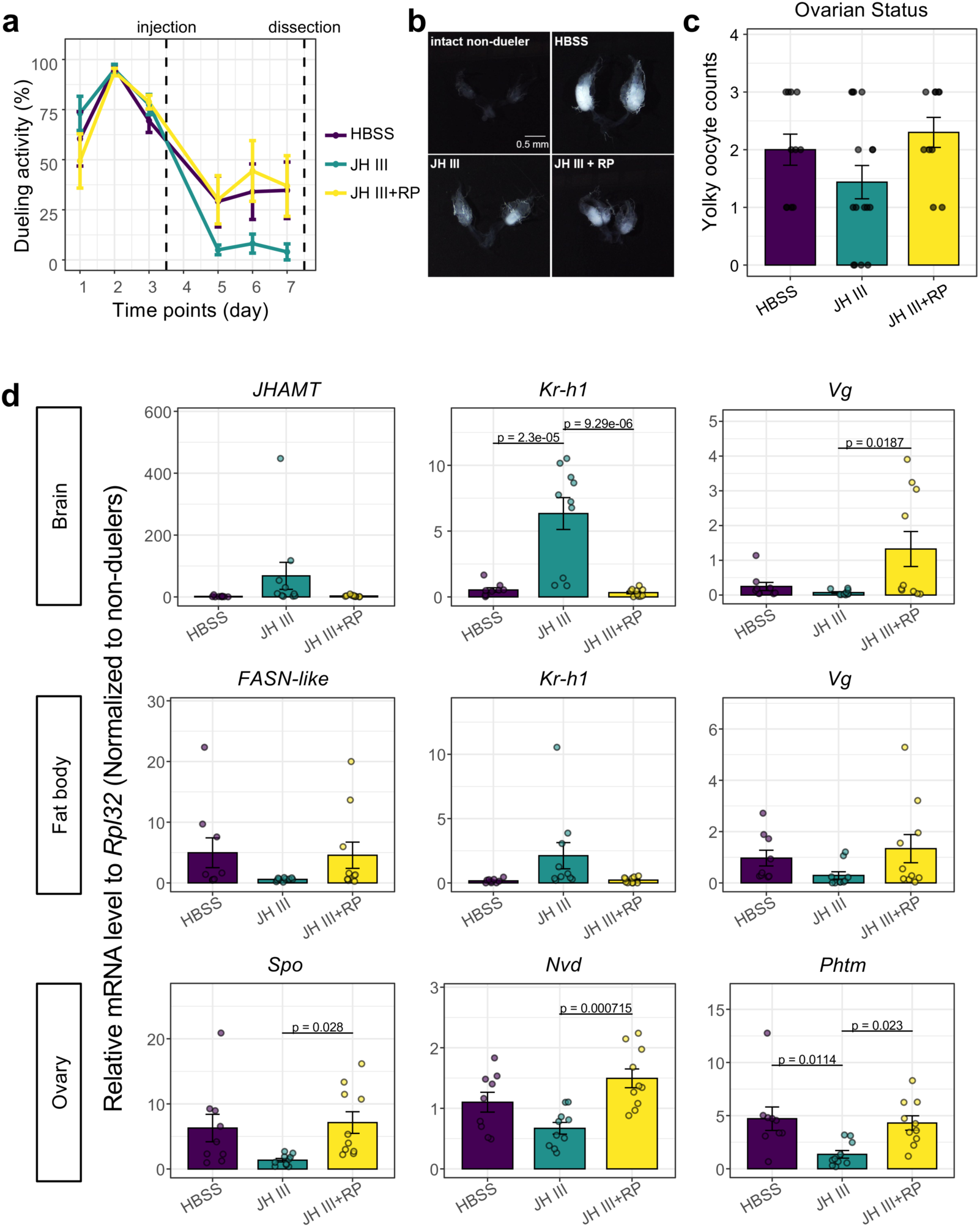
Recombinant HCRG1 reverses the effect of JH III. **a,** Quantification of dueling activity across treatment groups. Error bars represent SEM. *p*-values were determined by one-way ANOVA followed by Tukey’s post hoc analysis. **b,** Representative images of dissected ovaries collected four days post-injection. Scale bar, 0.5 mm. **c,** Quantification of yolky oocytes across treatment groups. Error bars represent SEM. **d,** qPCR analysis of gene expression in brains (top), fat bodies (middle), and ovaries (bottom) from different treatment groups. *RPL32* was used as an internal control. Results from multiple colonies were combined, and each expression value was normalized to the average of intact non-duelers from the same colony. Error bars represent SEM. *p*-values were determined by one-way ANOVA followed by Tukey’s post hoc analysis. Hank’s Balanced Salt Solution (HBSS) was used as a mock control. Sample sizes: HBSS (n = 9), JH III (n = 10), JH III + RP (n = 10).

To assess the molecular responses to treatment with JH III alone or in combination with HCRG1 RP, we examined the expression of key JH target genes across the brain, fat body, and ovary. In both the brain and fat body, *Kr-h1* was markedly upregulated following JH III treatment, but this induction was abolished in the presence of HCRG1 RP (**Fig. 5d**). In contrast, *Vg* was downregulated upon JH III injection, but restored to high levels under combined treatment with HCRG1 RP (**Fig. 5d**). Moreover, *JHAMT*, a key gene in JH biosynthesis and the most significantly upregulated gene in the brain following *HCRG1* KD (**Fig. 4b**), was induced by JH III, but this induction was neutralized by co-injection with HCRG1 RP (**Fig. 5d**). In the fat body, *FASN-like*, one of the most significantly downregulated genes upon *HCRG1* RNAi (**Fig. 4c**), was also suppressed by JH III, an effect that was reversed by HCRG1 RP. In the ovary, we assessed three representative Halloween genes—*Spo*, *Nvd*, and *Phtm*—which were expressed at higher levels in duelers than in non-duelers (**Fig. 5d**). JH III treatment led to a substantial suppression of their expression, and this repression was alleviated by HCRG1 RP (**Fig. 5d**).

Taken together, these results support the model that HCRG1 counteracts JH activity, thereby inducing the worker-to-gamergate transition.

### HCRG1 evolved as a lineage-specific regulator of conserved Hexamerin-mediated JH sequestration

Although JH suppresses reproductive caste transition in adult *Harpegnathos*, this negative relationship between JH and reproduction is not a universal feature of insects. In many solitary insects, including *Drosophila melanogaster*, JH functions as a gonadotropic hormone^42^. Moreover, JH regulation has been extensively rewired during the evolution of eusocial lineages, including bees and ants^16,17,43^. These observations raise the possibility that the HCRG1–Hex70c complex represents an evolutionarily derived mechanism for modulating JH physiology in social insects.

To examine this, we analyzed the molecular evolution of *HCRG1* and *Hex70c* across a curated 70-species insect dataset spanning diverse social traits (**Fig. 6a**). Tree-reconciliation-based orthology inference with OrthoFinder identified Hex70c co-orthologs in 66 species, whereas *HCRG1* co-orthologs were more restricted and absent from many sampled taxa, including *Drosophila* (**Fig. 6a**), indicating that *Hex70c* is broadly retained across insects, while *HCRG1* is more lineage-restricted. Codon-based terminal two-ratio branch scans further revealed contrasting patterns of branch-level ω evolution between the two genes. *HCRG1* showed terminal-branch elevations in ω in multiple ant species within Ponerinae, the *Harpegnathos*-containing clade that is broadly associated with gamergate reproductive systems, as well as in Ectatomminae, which includes a non-ponerine gamergate species (**Fig. 6a**). In contrast, *HCRG1* showed reduced foreground ω in Diptera and in the solitary carpenter bee *Xylocopa violacea*, consistent with stronger selective constraint in these lineages. *Hex70c* showed more broadly distributed ortholog retention and generally lower absolute ω values with more modest terminal-scan ω shifts; notably, the *Harpegnathos* terminal branch showed reduced *Hex70c* foreground ω, consistent with stronger constraint on this conserved hexamerin scaffold (**Fig. 6a**).

**Fig. 6.**
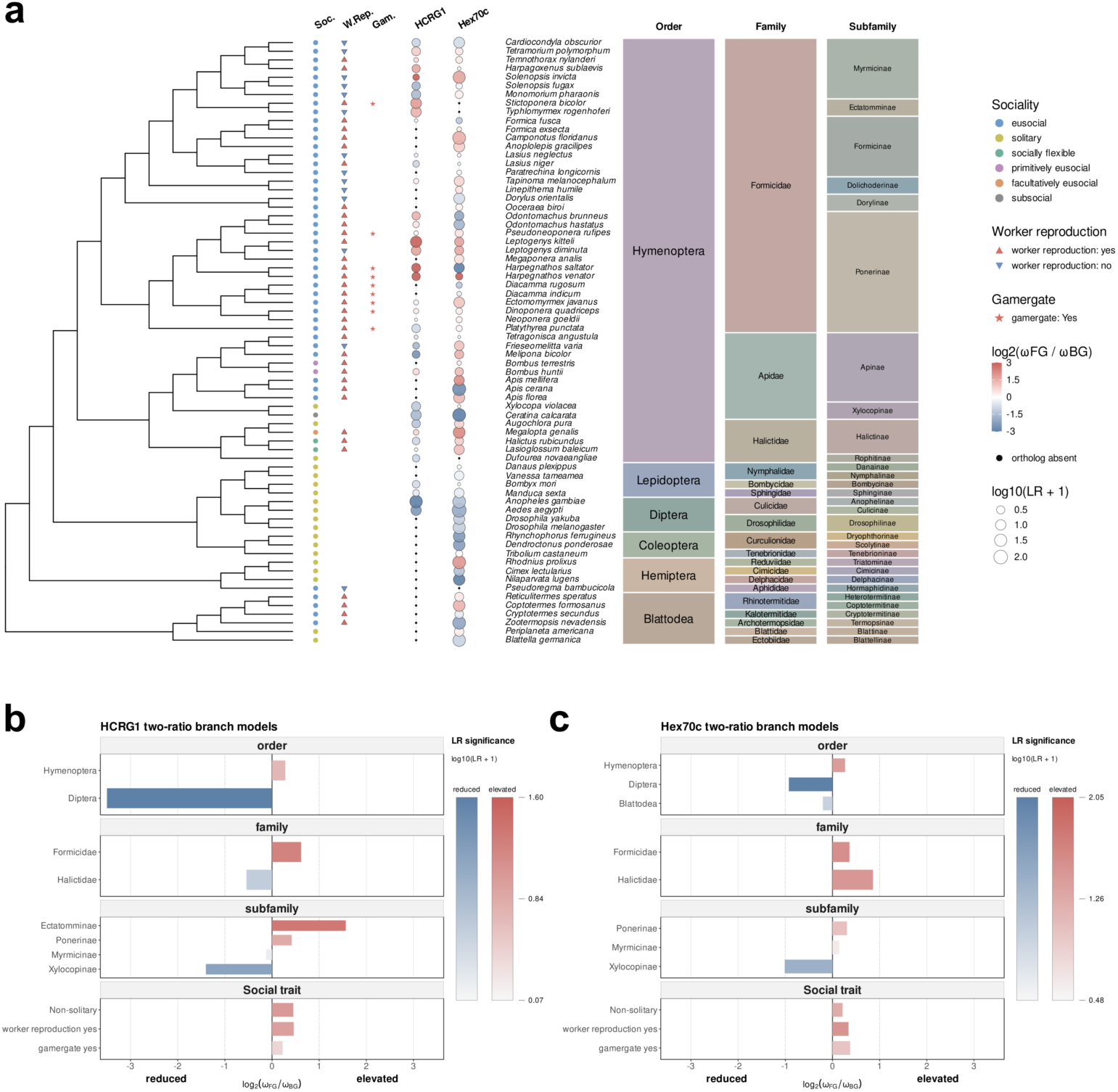
Phylogenetic distribution and clade-level branch-model signals of HCRG1 and Hex70c evolution. **a,** A 70-species insect cladogram annotated with sociality, worker reproductive capacity, gamergate status, terminal branch signals for HCRG1 and Hex70c, and taxonomic classification. Sociality is shown by colored circles, worker reproduction by filled triangles, and gamergate presence by red stars. HCRG1 and Hex70c terminal-scan results are displayed as bubble tracks, where color indicates log₂(ω_foreground/ω_background) from reduced foreground ω in blue to elevated foreground ω in red, and bubble size represents log₁₀(LR + 1). Black dots mark species lacking an ortholog and therefore lacking a terminal-scan result. Taxonomic order, family, and subfamily are shown as colored blocks to the right of the species labels. **b,c,** Centered bar plots show foreground-specific shifts in ω for HCRG1 (b) and Hex70c (c) two-ratio branch models. Bars extending left indicate reduced foreground ω, whereas bars extending right indicate elevated foreground ω, with bar extension along the x axis representing log₂(ω_FG/ω_BG). Foregrounds are grouped by taxonomic level and social trait. Blue and red bars indicate reduced and elevated foreground ω, respectively, and color intensity represents likelihood-ratio support, shown as log₁₀(LR + 1), with darker colors indicating stronger support within each direction.

Foreground-based two-ratio branch models further supported lineage- and trait-associated ω differences in *HCRG1*. *HCRG1* showed elevated foreground ω in several ant and social foregrounds, including Formicidae, Ponerinae, non-solitary lineages, and lineages with worker reproductive capacity, whereas reduced foreground ω was observed in Diptera and Xylocopinae, consistent with stronger constraint in these lineages (**Fig. 6b**). The strongest *HCRG1* branch-model signal was taxonomic rather than strictly trait-specific: Ectatomminae showed the largest elevation and exceeded ω = 1, while Ponerinae and Formicidae showed more moderate foreground ω elevations. Multi-ratio branch models resolved a similar taxonomic pattern, with elevated class-specific ω in Ectatomminae and Ponerinae, and with Formicidae elevated relative to other Hymenopteran families (**Extended Data Fig. 8a**). Trait-based multi-ratio models provided supportive but less specific evidence: *HCRG1* ω was elevated in eusocial or facultatively eusocial classes and in worker-reproductive lineages, whereas within Formicidae, worker-reproductive and non-worker-reproductive classes had similar ω values, and gamergate status did not correspond to the strongest ω elevation. By contrast, *Hex70c* showed generally low absolute branch-level ω values and less consistent foreground-to-background differences. Two-ratio models detected elevated foreground ω in several social or ant-associated foregrounds, including Formicidae, Ponerinae, worker-reproductive lineages, and gamergate-associated lineages, but these elevations remained modest in absolute magnitude compared with *HCRG1* (**Fig. 6c**). Multi-ratio models further resolved class-specific heterogeneity in Hex70c, including elevated ω in some Hymenopteran, Formicidae, and termite-associated classes, while other classes showed reduced or near-background ω (**Extended Data Fig. 8b**). Together, these results indicate that *HCRG1* underwent repeated Formicidae-centered elevations in branch-level ω, consistent with accelerated protein evolution and reduced selective constraint and/or adaptive change, whereas *Hex70c* retained generally low ω values consistent with stronger functional constraint on a conserved hexamerin scaffold. Consistent with this model, HCRG1 residues identified by PAML BEB analysis in global site and branch-site models mapped close to the HCRG1–Hex70c interface in the structural model (**Extended Data Fig. 9**), supporting a model in which lineage-specific evolution of *HCRG1* contributed to the emergence or refinement of its interaction with a pre-existing Hexamerin scaffold.

If HCRG1 is able to bind a conserved Hexamerin ortholog, introduction of ant HCRG1 into *Drosophila*, where HCRG1 orthologs are absent, would be expected to artificially generate an HCRG1–Hexamerin complex. In the OrthoFinder results, we identified DmLsp2 as the closest *Drosophila* ortholog of ant Hex70c. AlphaFold3 modeling of ant HCRG1 with DmLsp2 yielded a high-confidence predicted complex (ipTM = 0.88, pTM = 0.90), suggesting that HCRG1 can potentially bind the conserved fly Hexamerin scaffold. We therefore compared ligand docking to DmLsp2 alone and HCRG1-bound DmLsp2. HCRG1 binding significantly increased the predicted affinity of DmLsp2 not only for JH III, but also for JH III bisepoxide (JHB3) and methyl farnesoate (MF) (**Extended Data Fig. 7c**), two major endogenous JH-related sesquiterpenoids in *Drosophila* ^44,45^. Together, these results support a model in which ant HCRG1 evolved as a lineage-specific modulator of a conserved Hexamerin protein, thereby creating an HCRG1–Hexamerin complex that may antagonize JH signaling through enhanced Hexamerin-mediated ligand sequestration.

### Ectopic expression of ant HCRG1 extends lifespan in *Drosophila*

Having established that ant HCRG1 can potentially engage the conserved fly Hexamerin Lsp2 and enhance its predicted association with JH-related ligands, we next asked whether ectopic expression of *Harpegnathos HCRG1* could produce a physiological outcome consistent with reduced JH signaling. Beyond their social and reproductive differences from workers, gamergates exhibit a dramatic extension of lifespan, from approximately seven months in workers to nearly three years in gamergates^5,6^. Since JH deprivation in adult insects is widely associated with increased longevity^22,46^, and we have shown that HCRG1 is both a JH antagonist and a positive regulator of the worker-to-gamergate transition, we hypothesized that HCRG1 might also contribute to lifespan extension. HCRG1 may exert long-term effects on gamergate lifespan due to its elevated expression in both duelers and mature gamergates^2^.

To test this possibility, we used female *Drosophila* as a model given its short lifespan that facilitates longevity assays that could not be performed in ants within a reasonable timeframe. We cloned *Harpegnathos HCRG1* and placed it under *UAS* control to perform tissue-specific expression via Gal4 drivers. Given that the brain and fat body are major sources of HCRG1^2^, we used *elav-Gal4* (pan-neuronal), *repo-Gal4* (pan-glial), *ppl-Gal4* and *Cg-Gal4* (abdominal fat body), and *3.1Lsp2-Gal4* (head fat body) for targeted expression. As a control, we also expressed *UAS-dFOXO*, a well-characterized longevity-promoting factor. Consistent with previous reports^47–49^, neuronal overexpression of *dFOXO* reduced median lifespan (**Fig. 7a**), whereas overexpression in glial and fat tissues extended it by 8–12 days compared to controls (**Fig. 7b-e**). *repo*-Gal4-driven *HCRG1* expression in glial cells significantly prolonged lifespan, increasing the median survival by 11 days compared to *EGFP* controls, comparable to *dFOXO* (**Fig. 7b**), while its expression in the abdominal fat body only resulted in a modest increase (**Fig. 7c and d**). Since single-cell sequencing data showed that *HCRG1* expression is higher in a specific glial subclass in duelers^12^ and that hemolymph HCRG1 and its effects on gamergate traits originate primarily from the dueling brain (**Fig. 2 and Extended Data Fig. 2b**), the increased longevity from glial *HCRG1* overexpression is consistent with this finding. Ectopic expression of ant *HCRG1* in neurons or the head fat body led to reduced lifespan, with median survival decreasing from 66 to 60 days and from 61 to 56 days, respectively (**Fig. 7a and e**), possibly due to excess HCRG1 leading to detrimental effects by perturbing secretory activity in these tissues.

**Fig. 7.**
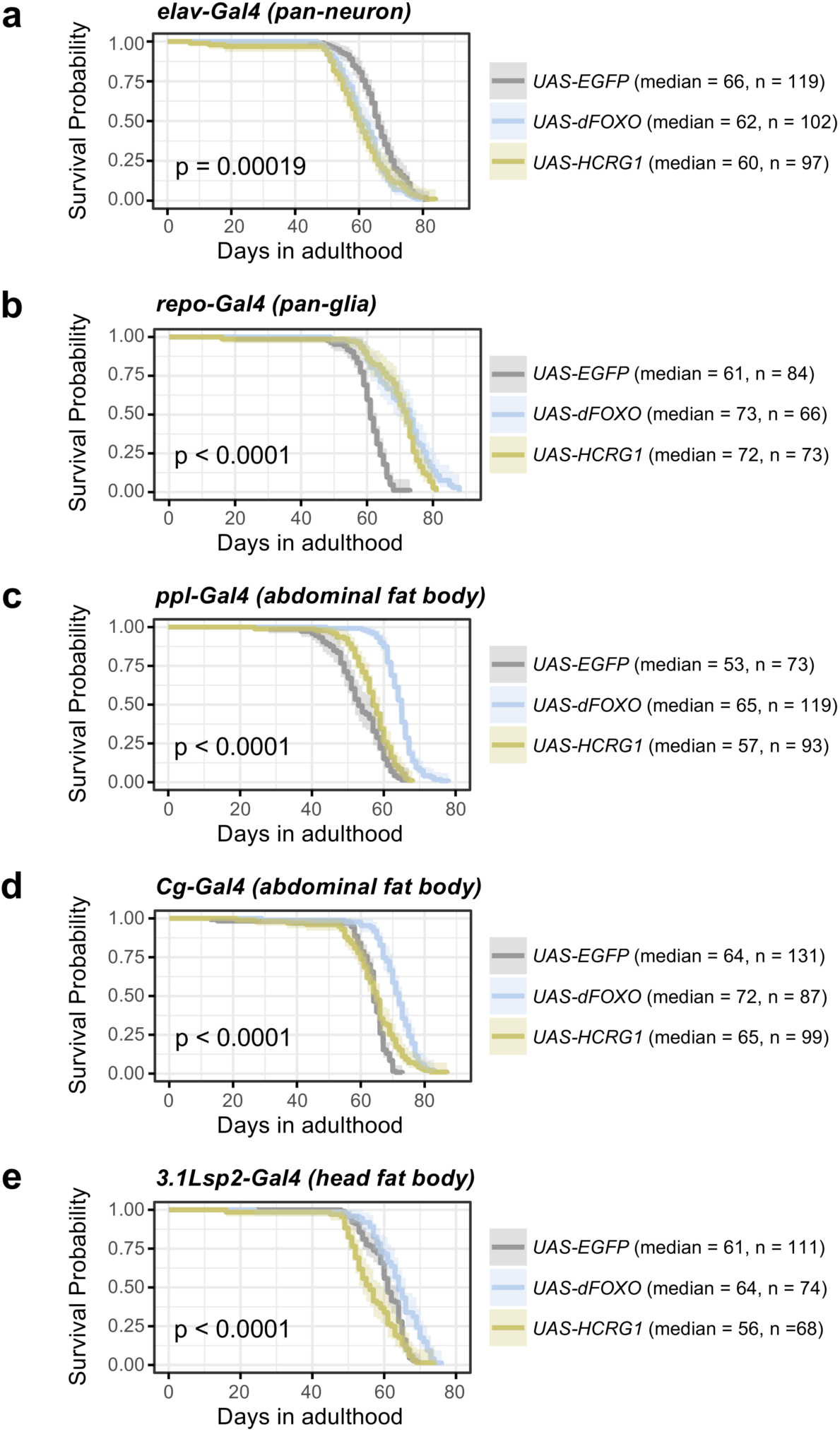
Ectopic HCRG1 affects lifespan in *Drosophila*. **a-e,** Survival curves of flies expressing *EGFP*, *dFOXO*, or ant *HCRG1* under the control of *elav*-Gal4 (**a**), *repo*-Gal4 (**b**), *ppl*-Gal4 (**c**), *Cg*-Gal4 (**d**), and *3.1Lsp2*-Gal4 (**e**). All Gal4 driver lines were combined with *tub-Gal80ts* for temporal control of expression. The number of recorded death events and median survival values are indicated. *p*-values were calculated using the log-rank test.

Taken together, our results suggest that HCRG1 supplementation mimics the lifespan-extending effects of JH deprivation in female *Drosophila* ^20–22^ (**Extended Data Fig. 7c**), thus supporting a potential role in promoting the extended gamergate longevity.

## DISCUSSION

Through integrated proteomic, structural, evolutionary, and functional analyses, we have identified HCRG1 as a lineage-specific suppressor of JH signaling that promotes the worker-to-gamergate transition in *Harpegnathos* ants, likely acting downstream of brain changes initiated by the loss of queen pheromones.

### Linking brain responses to organism-wide plasticity

In *Harpegnathos*, worker reproduction is suppressed by queen pheromones, which are detected by olfactory receptors in the antennal sensilla^8,9,50^ and subsequently alter brain gene expression^2,6,10,12,14^. In the presence of queen pheromones, the worker brain maintains high JH activity, leading to the expression of *Kr-h1*^14^ and *Crz* to sustain the worker fate^10^. Upon queen loss, some workers become gamergates by downregulating JH and upregulating 20E, accompanied by increased brain insulin to support the development of the ovaries^6^.

Through our profiling of the hemolymph of transitioning ants, we have shown that circulatory HCRG1 levels increased in gamergate candidates (duelers), likely supported by its increased expression in the brain of early duelers^2^. This increase in circulatory HCRG1 is required for dueling behavior (**Fig. 2d**), vitellogenesis in abdominal fat body (**Fig. 4c**), and ovarian development (**Fig. 2e-f and 4d**). We found that neither RNAi-mediated KD of *HCRG1* nor ectopic treatment with HCRG1 RP affected *Crz* expression in the brain. However, *Crz receptor* (*CrzR*) KD was previously shown to increase *HCRG1* expression in the central brain^10^. Together with the observed reduction in *Crz* levels in duelers (Extended Data Fig. 2d), these findings suggest a possible asymmetric relationship between Crz signaling and HCRG1 regulation. If *Crz* and *HCRG1* are functionally connected, *Crz* signaling is more likely to act upstream of *HCRG1*, potentially as a suppressive input that is relieved during dueling. Increased HCRG1 would then mediate brain-derived signals for systemic reprogramming during caste differentiation. This model remains speculative, as the relevant *CrzR*-expressing cells and their relationship to queen-pheromone-responsive circuits remain unresolved.

### Kunitz domain-containing proteins and reproductive development

HCRG1 contains a highly conserved Kunitz domain^51,52^, which is typically found in protease inhibitors. Knock-down of *HCRG1* in duelers suppressed ovarian development, whereas injection of recombinant HCRG1 protein in non-duelers promoted it (**Fig. 2e-f and 3c-d**). This suggests that HCRG1 functions as a regulator of reproductive maturation. In mammals, Kunitz domain-containing proteins have also been implicated in female reproductive development^53^. For example, in humans, reduced protein levels of SPINT1 (a homolog of HCRG1) in maternal plasma are associated with placental insufficiency and adverse pregnancy outcomes^54,55^. In mice, SPINT1 knockout leads to matriptase-dependent placental failure^56^, defective placental labyrinth formation, and abnormalities in embryonic neural tube closure in mice^57^. Our discovery that HCRG1 modulates ovarian development and caste differentiation in *Harpegnathos* supports an ancient role for Kunitz domain proteins in reproductive plasticity and developmental remodeling, although in this case, HCRG1 appears to act not as a protease inhibitor, but through its binding to Hexamerin.

### Regulation of JH and its evolutionary implications

In solitary insects, reproductive development and lifespan are typically negatively correlated^58^. In these species, JH promotes both reproductive maturation and a short lifespan. JH controls ovarian development by binding to its receptor Methoprene-tolerant, which acts through Kr-h1 to regulate the expression of genes essential for vitellogenesis and oocyte development^42,59–63^. However, reducing JH production, either through removal of the corpora allata (the JH-producing gland) or by down-regulating insulin, suppresses reproduction and extends lifespan^20,22,64^.

In social insects, the interplay between JH and reproduction is more complex: in ants and many eusocial hymenopterans, JH has lost its ancestral gonadotropic role and regulates the worker caste^32,65^, repressing reproduction by antagonizing the action of 20E^2,14,33,66,67^, which is required for ovarian development^37–41^. This decoupling of JH from reproduction, particularly vitellogenesis in the worker caste, is a key focus of the reproductive ground plan hypothesis, which posits that social behaviors evolved from ancestral reproductive pathways^32,68–70^. The retention of the role of JH in regulating some reproductive behaviors like aggression and brood care in worker castes suggests that JH signaling has been co-opted during the evolution of eusociality, a process in which the JH pathway has undergone recurrent positive selection^17^. As a consequence, the few reproductive individuals (queens) must have lower JH titers than non-reproductives^2,14,66^, and they also exhibit greater longevity^71^. This shift implies that regulation of the JH pathway is under strong evolutionary pressure. Adding another layer of regulation, JH-binding proteins such as Hexamerins appear to have undergone parallel evolutionary adaptations. Hexamerins exhibit convergent and complementary signatures of selection associated with social gains and losses^16^. In Hymenoptera, Hexamerins are classified into two groups based on molecular weight: high-molecular-weight Hexamerin 1 (∼110 kDa) and low-molecular-weight Hexamerin 2 (∼70 kDa)^72^. Direct binding of JH by Hexamerins has been demonstrated in Orthoptera and termites: in locusts, ∼70 kDa hexameric storage proteins show a high affinity for JH III, the primary JH type in ants^73,74^; in termites, the JH-binding Hexamerin is approximately 85 kDa^19^. These reported molecular weights are similar to that of Hex70c, which we identified as an HCRG1 interactor in *Harpegnathos*, further supporting a conserved role for this subclass of Hexamerins in JH binding. Consistent with this view, our comparative analysis showed that Hex70c co-orthologs are broadly retained across insects, whereas HCRG1 co-orthologs are much more restricted and are absent from many sampled taxa (**Fig. 6 and Extended Data Fig. 8**), including *Drosophila*. This asymmetry suggests that the HCRG1–Hex70c complex did not arise through parallel emergence of both components, but instead through the evolution of a lineage-specific HCRG1 factor capable of engaging a pre-existing conserved Hexamerin scaffold. The enrichment of positively selected HCRG1 sites near the HCRG1–Hex70c interface further supports the idea that selection acted on HCRG1 to create or refine this interaction (**Extended Data Fig. 9**). Thus, HCRG1 may represent an evolutionarily derived regulatory layer superimposed on an older Hexamerin-based JH-binding system.

In *Harpegnathos*, JH III injection not only suppresses egg production but also promotes non-reproductive social behaviors^2,14^. In gamergates, circulating HCRG1 levels are elevated and are necessary for promoting both ovarian development and dueling behavior by acting as an antagonist of JH activity. Indeed, treatment with HCRG1 RP can counteract the inhibitory effects of JH III on ovarian development and dueling activity. JH III is a highly lipophilic molecule that requires carrier proteins for transport in the aqueous hemolymph^23^. The interaction between HCRG1 and Hex70c provides a possible mechanism by which HCRG1 regulates JH titers. Circulating Hex70c protein levels do not differ between duelers and non-duelers (**Extended Data Fig. 1a**). Our data support a model in which HCRG1 regulates the functional state of a pre-existing Hex70c pool. In termites, loss-of-function Hexamerin leads to elevated JH titers, which in turn promotes the differentiation of workers into the pre-soldier stage^26^. However, it is not known whether Hexamerin alone can suppress JH. Structural modeling and JH III docking support a mechanism in which HCRG1 enhances the JH-binding affinity of Hex70c (**Extended Data Fig. 7a**). The ability of ant HCRG1 to form a predicted complex with *Drosophila* Lsp2 and increase the predicted association of Lsp2 with JH-related ligands further supports the idea that HCRG1 acts by modifying a conserved Hexamerin scaffold. We hypothesize that Hex70c functions as a “cage” for JH, and HCRG1 as a “lock” that secures JH within this cage, stabilizing the complex and reducing JH bioavailability (**Extended Data Fig. 10**). In non-duelers, despite the presence of sufficient Hexamerin “cages” in the hemolymph, the absence of enough “locks” (HCRG1) prevents JH from being sequestered. In duelers, increased HCRG1 production enables the formation of a stable HCRG1-Hexamerin complex, effectively trapping JH and reducing its bioavailability, thereby promoting gamergate differentiation.

The observation that HCRG1 overexpression extends lifespan in *Drosophila* further underscores the conserved role of JH suppression in promoting longevity across insects (**Fig. 7b**). Since *Drosophila* lacks an endogenous HCRG1 ortholog, this experiment also provides a heterologous test of whether ant HCRG1 can impose an ant-like JH-antagonistic module onto a conserved fly Hexamerin system. In many eusocial species, including insects, birds, and mammals, reproductive individuals often live significantly longer than non-reproductive members of the colony^75–79^. This pattern contrasts with the typical fecundity-longevity trade-off observed in solitary organisms. In eusocial insects, JH not only suppresses reproduction but also promotes aging^33,80,81^, making its downregulation essential for achieving caste-specific longevity. In our previous work, we showed that reproductive *Harpegnathos* ants overcome the pro-aging effects of insulin signaling^6^, providing a necessary mechanism for extended lifespan. Here, we identify HCRG1-mediated JH antagonism as a complementary mechanism that may help explain how reproductive ants decouple fecundity from longevity. By modulating a conserved Hexamerin scaffold, HCRG1 may reduce JH bioavailability while permitting the reproductive and metabolic programs required for gamergate maturation.

## Acknowledgements

We thank all members of the Desplan and Reinberg laboratories for helpful discussions and advice. We are grateful to Dr. Sanxiong Liu and J. Granat for their insightful suggestions and technical assistance. We thank Dr. Samuel Church for constructive advice on the evolutionary analyses. We thank Drs. Richard Binari, Marc Freeman, Daniela Drummond-Barbosa, Heinrich Jasper, and Gilles Storelli for providing the Gal80ts combined Gal4 fly strains *elav-Gal4*, *repo-Gal4*, *3.1Lsp2-Gal4*, *ppl-Gal4*, and *Cg-Gal4*, respectively. This work was supported in part through the NYU IT High Performance Computing resources, services, and staff expertise.

## Author contributions

L.D. designed and performed the majority of experiments under the supervision of C.D. and D.R. J.M. conducted most of the behavioral analyses, assisted with ant source maintenance, and contributed to molecular and biochemical assays. H.Z. performed all mass spectrometry analyses. C.L. contributed to the fly survival assays. Y.C. conducted the upstream bioinformatic analysis of TMT data. L.D. and C.D. wrote the manuscript with input from the other authors.

## Data and materials availability

The RNA-seq data generated in this study have been deposited in NCBI’s Gene Expression Omnibus (GEO) under accession number GSE298046. The data will be publicly available upon publication or on December 30, 2025, whichever comes first. The mass spectrometry proteomics data have been deposited to the ProteomeXchange Consortium via the MassIVE partner repository under the dataset identifier MSV000098013. The data are currently under embargo and will be made publicly available upon publication.

## Materials and Methods

### *Harpegnathos* ant rearing and transition colony setup

*Harpegnathos saltator* colonies were originally obtained from Jürgen Liebig’s laboratory at Arizona State University. As previously described ^2,6^, ants were maintained in plastic boxes (Pioneer Plastics, Inc.) containing an underground arena covered with glass slides (Darby Dental, 849-1560), and kept in a temperature-controlled room at 25 ± 2 °C under a 12-hour light–dark cycle. Colonies were provided with live crickets three times per week.

For the transition colony setup, 30 young female adults (2 weeks post-eclosion) were uniquely labeled with Uni-Paint markers and introduced into plastic boxes as described above. Transition colonies were fed pre-stung crickets with the same frequency as source colonies. To minimize feeding-related variability, all colonies were starved for 24 hours before sampling. Individuals that died prior to sampling were excluded from the analysis.

### Hemolymph extraction

Ants were first rinsed in distilled water, and all six legs and gasters were carefully removed using Vannas scissors (WPI, 501232). The remaining head and thorax were transferred into a 0.5 mL microcentrifuge tube with a small hole at the bottom, created using a 23-3/4 gauge needle. This tube was placed inside a 1.5 mL microcentrifuge tube, and hemolymph was collected by centrifugation at 3,000 × g for 1 minute at 4 °C.

### TMT proteomic analysis

Hemolymph samples were collected from duelers and non-duelers at three time points (3, 10, and 30 days) after the onset of dueling, with four biological replicates per condition. Additionally, four pre-dueling individuals (day 0) were included as a baseline control. For each sample, 50 µg of protein was prepared along with a pooled reference sample (1:1 mix of all samples). Samples were separated using NuPAGE 10% Bis-Tris Gel (1.5 mm × 10 well, Invitrogen) as gel plugs. The gel was stained with Coomassie Brilliant Blue R250, destained, and subjected to in-gel digestion using a standard protocol.

The digested peptides were divided into two sets, each including two bridge samples (identical between sets) for downstream inter-batch correction. Samples in each set were labeled with Thermo TMTpro (16-plex, Lot #: UI2192951) following the manufacturer’s instructions. After labeling, the samples were pooled, dried under vacuum, desalted using Sep-Pak tC18 cartridges (Varian, WAT054960), and fractionated by high-pH reverse-phase liquid chromatography (RPLC) using an Agilent 1100 series system. Peptides were injected onto an Xbridge column (Waters, C18, 3.5 µm, 2.1 × 150 mm) and separated using a linear gradient of 1% B per minute from 2% to 45% of buffer B (buffer A: 20 mM ammonium, pH 10; buffer B: 20 mM ammonium in 90% acetonitrile, pH 10). UV absorbance at 214 nm was monitored, and 1-minute fractions were collected. Selected fractions were desalted using StageTips and analyzed by nano-LC-MS/MS.

Nano-LC-MS/MS was performed using a Dionex rapid-separation liquid chromatography system coupled to an Eclipse mass spectrometer (Thermo Fisher Scientific). Samples were first loaded onto an Acclaim PepMap 100 trap column (75 µm × 2 cm, Thermo Fisher) and washed with Buffer A (0.1% trifluoroacetic acid) for 5 minutes at a flow rate of 5 µl/min. The trap column was then brought in line with the analytical column (nanoEase, MZ Peptide BEH C18, 130Å, 1.7 µm, 75 µm × 20 cm, Waters) at a flow rate of 300 nL/min using a multistep gradient: 4%–15% buffer B (0.16% formic acid and 80% acetonitrile) over 20 minutes, 15%–25% buffer B over 40 minutes, and 25%–50% buffer B over 30 minutes.

The MS acquisition began with an MS1 scan (Orbitrap analysis, resolution 120,000, scan range from 350 to 1600 m/z, automatic gain control [AGC] target 1E6, maximum injection time 100 ms). The top-speed mode (3-second duty cycle) was used to determine the number of MS/MS scans per cycle. Parent ions were isolated using the quadrupole (isolation window 1.2 m/z, AGC target 1E5) and fragmented with higher-energy collisional dissociation (normalized collision energy 33%). Fragment ions were detected in the Orbitrap (resolution 30,000). The MS/MS scan range was determined by the charge state of the parent ion, with a lower limit of 110 m/z.

LC-MS/MS data were processed using Proteome Discoverer 2.4 (Thermo Fisher Scientific) with the Sequest HT search engine against the *Harpegnathos* saltator protein database (Hsal_v8.5, GenBank assembly accession GCA_003227715.1) and a common contaminant database. MS mass tolerance was set to ±10 ppm, and MS/MS mass tolerance was set to ±0.02 Da. TMTpro labeling on lysine residues and peptide N-termini, as well as carbamidomethylation on cysteine, were set as static modifications. Methionine oxidation, protein N-terminal acetylation, N-terminal methionine loss, and N-terminal methionine loss plus acetylation were set as dynamic modifications. Peptide identification was validated using Percolator, and a target-decoy strategy was employed to estimate the false discovery rate (FDR), with high-confidence peptides defined at FDR <0.01 and medium-confidence peptides at FDR <0.05.

For quantification, reporter ion abundance was calculated using the signal-to-noise ratio (S/N) if available; otherwise, intensity values were used. Reporter ion intensities were corrected for isotopic impurities, with a co-isolation threshold set at 50%, average reporter S/N threshold at 10, and SPS mass matches percentage threshold at 65%. Protein abundance in each channel was calculated using the summed S/N of all unique and razor peptides and normalized to the total peptide abundance within each channel.

To correct for batch effects, common pooled internal standards were used to normalize reporter ion intensities between the two TMT experiments. After internal reference scaling (IRS) normalization ^82^, differential protein abundance between groups was determined by comparing IRS-normalized total reporter ion intensities using the Bioconductor packages dplyr and limma. The MPI Bioinformatics Toolkit (https://toolkit.tuebingen.mpg.de/tools/hhpred) was employed to identify potential homologs of uncharacterized components through HHPred-based profile–profile comparisons.

### RNA isolation and qPCR

Tissues from individual *Harpegnathos* ants were dissected and homogenized using an RNase-free pellet pestle (Fisher, Cat# 12-141-368) in a 1.5 mL microcentrifuge tube containing 200 µL of TriPure reagent (Roche, Cat# 11667157001). RNA was extracted using phenol-chloroform extraction and ethanol precipitation. Genomic DNA was removed, and reverse transcription was performed using the QuantiTect Reverse Transcription Kit (Qiagen, Cat# 205314) according to the manufacturer’s instructions. mRNA expression levels were measured by quantitative PCR (qPCR) using SYBR Green I Master Mix (Roche, Cat# 04707516001) or Luna Universal qPCR Master Mix (NEB, Cat# M3003S) on a Bio-Rad CFX96 real-time PCR system.

### HCRG1 recombinant protein and custom antibody

The construct design is shown in Figure 3A. The fragment was cloned into the pFastBac vector, and the full-length recombinant protein was expressed in Sf9 cell cultures according to the Bac-to-Bac® Baculovirus Expression System handbook (Invitrogen). The expressed recombinant protein was purified using Ni-NTA Agarose beads (QIAGEN, Cat# 30210) and eluted with 100 mM imidazole. The elution was then dialyzed against 1×PBS to exchange the buffer. A portion of the protein in PBS was used for generating polyclonal antibodies against HCRG1. The epitope used for rabbit immunization was the full-length recombinant HCRG1 protein with the following sequence:

MGLKSCLLFTLIIVGILSGSSGSSHHHHHHGSSGSSENLYFQGHEIVAKPSSICQLPKVVGPCRASLKRYRYDSTTGQCEEFTYGGCKGNENNFITREVCQENCINN

Another aliquot of the purified protein in PBS was incubated with TEV enzyme. After cleavage efficiency was confirmed via Tris-Tricine gel analysis, the remaining fragments, which did not contain the His tag, were recovered using a column and dialyzed into Hank’s Balanced Salt Solution (HBSS) for use in RP treatments.

### Ponceau stain and western blots

Hemolymph was extracted and homogenized in lysis buffer (50 mM Tris-HCl, pH 7.8, 150 mM NaCl, and 1% Nonidet P-40). The lysate was mixed with 2× Tris/Tricine loading buffer and boiled at 100 °C for 5 minutes. Boiled samples were loaded onto Tris/Tricine/SDS gels and transferred to nitrocellulose membranes. Membranes were stained immediately after transfer using Ponceau S Staining Solution (Thermo Scientific, Cat# A40000279), followed by several washes with distilled water until the background was fully removed. Stained membranes were imaged using a standard scanner, and band intensity was quantified using Fiji.

The Ponceau dye was washed away by blocking the membranes in TBST buffer (150 mM NaCl, 20 mM Tris-HCl, pH 7.5, 0.05% [v/v] Tween-20) containing 3% non-fat dry milk (Carnation) for 1 hour at room temperature. Membranes were then incubated with primary rabbit anti-HCRG1 antibody (1:1000) diluted in TBST with 3% milk overnight at 4 °C. Anti-rabbit IgG HRP-conjugated secondary antibody (Promega, Cat# W401B) was used at a 1:5000 dilution in TBST with 3% milk. Bands were detected using West Femto Maximum Sensitivity Substrate (Thermo Fisher, Cat# 34094), and band intensity was quantified using Fiji.

### Behavior monitoring and measurement

Experimental colonies were set up under recording cameras (Logitech C920 webcam), and videos of the inside of the nests were recorded using the method described previously^2^. Dueling activity was scored every 45 minutes during the 12-hour light cycle (a total of 16 times per day). The scoring method was based on the presence or absence of dueling behavior within the first 20 minutes of each time window.

### Injection experiments

Custom dicer-substrate short interfering RNAs (DsiRNAs) targeting *HCRG1* and *GFP* were synthesized by IDT and resuspended in RNase-free water. Equal volumes of DsiRNAs and a 10% glucose solution were mixed to a total volume of 10 μL. Separately, 0.8 μL of in vivo-jetPEI reagent (Polyplus Transfection) was diluted in 10 μL of glucose solution (final concentration of 5%). The two solutions were combined, yielding working concentrations of DsiRNAs (2.5 μM, 5 μM, and 10 μM for HCRG1 and 10 μM for GFP). The mixture was incubated for 15 minutes at room temperature before injection. Each ant was injected with 0.5 μL of the DsiRNA-PEI complex directly into the head through a hole made between the antennal stems using a steel pin (FST 26002-15) and a calibrated glass capillary (Sutter Instrument, Item# Q100-70-7.5) pulled using a micropipette puller (Sutter Instrument, model P-2000, program 46).

For RP rescue treatments, an additional abdominal injection was performed alongside the head RNAi injection. A total of 1 μL of recombinant protein (100 ng/μL) in HBSS or blank HBSS (mock control) was injected into the abdomen through an opening between the tergites using a calibrated glass capillary under the same parameters as the head injection. For RP-treated non-duelers, day 3 non-duelers received a single abdominal injection following the same procedure used in the RP rescue experiments.

For JH III treatment, 10 mg of synthetic JH III (Sigma J2000) was dissolved in pure ethanol (EtOH) to achieve a concentration of 5 μg/μl. A 1 μl solution of HBSS containing 10% EtOH, 0.5 μg/μl JH III in HBSS containing 10% EtOH, or 0.5 μg/μl JH III combined with 100 ng/μl RP in HBSS containing 10% EtOH was injected into the head in the same manner as the RNAi assays.

Experimental colonies were monitored by cameras after setup. Injections were performed on duelers or non-duelers 3 days after the colony began dueling. Video recordings resumed 24 hours post-injections, and tissue samples were collected 3 days after injection.

### Ovary phenotyping

Ovary images were captured using a Nikon Stereomicroscope System with a camera (serial #1002455, #1008177, #10) or an Eyepiece Mount Swift 5.0 Megapixel Digital Camera. For each ovary, all eight ovarioles were examined, and only the most mature egg chambers from each ovariole were included in the final quantification. Individuals that died prior to sample collection were excluded from the analysis.

### Immunoprecipitation-mass spectrometry (IP-MS)

IP-MS was performed using HCRG1 antibody or rabbit IgG (control) crosslinked to Protein G agarose beads (Sigma, Cat# 11243233001).

#### Antibody crosslinking

Protein G agarose beads (200 µL) were washed three times with PBS by centrifugation at 500g for 1 minute. The beads were incubated with 20 µg of HCRG1 antibody or rabbit IgG in 1 mL PBS at room temperature for 2 hours with rotation. After incubation, the beads were washed twice with 10 volumes of 0.2 M sodium borate (pH 9.0) and resuspended in the same buffer containing 20 mM dimethyl pimelimidate (Thermo Scientific, Cat# 21667). The mixture was rotated at room temperature for 1 hour. The reaction was quenched by washing the beads once with 0.2 M ethanolamine (pH 8.0), followed by resuspension and incubation in the same buffer for 2 hours at room temperature with gentle mixing. The beads were washed with 0.2 M ethanolamine (pH 8.0), followed by a brief wash with 0.1 M glycine (pH 2.2) to remove loosely bound antibodies. Antibodies were refolded by washing with 1 M Tris (pH 8.0). Finally, the beads were washed three times with PBS and stored in 1 mL PBS containing 0.01% NaN₃.

#### Sample preparation

Whole ants were washed with distilled water to remove surface debris, dried on Kimwipes, and placed in pairs into 2 mL microcentrifuge tubes containing a single 7 mm stainless steel bead (Qiagen, Cat# 69990). Samples were snap-frozen in liquid nitrogen, transferred to a pre-cooled (-80 °C) tube holder, and homogenized for 120 seconds at 50 Hz using a TissueLyser LT (Qiagen, Cat# 85600). The homogenates were immediately placed on ice, and 1 mL of 1% digitonin lysis buffer (1% digitonin [Sigma, Cat# 300410], 50 mM HEPES [pH 7.2], 10 mM EDTA, 150 mM NaCl, 10 mM NaF, 1 mM PMSF, and protease inhibitor cocktail [Sigma]) was added. Lysates were incubated on ice for 2 hours and centrifuged at 12,000g for 10 minutes at 4 °C. The supernatants were pooled and dialyzed into PBS to equilibrate with the crosslinked beads for immunoprecipitation. Lysates were prepared from two independent colonies (initiated with 30 callow female ants), providing a total of 10 day-10 duelers and 10 day-10 non-duelers.

#### Immunoprecipitation

After dialysis, lysates from dueler and non-dueler groups were each used in duplicate for IgG control and anti-HCRG1 IP. Each 2 mL lysate was incubated with 200 µL of Protein G beads at 4 °C overnight with gentle rotation. Beads were washed three times with Wash Buffer (20 mM HEPES, 150 mM NaCl, 1.5 mM MgCl₂, 0.2 mM EDTA, and 5% glycerol).

#### Protein digestion

Two IP batches were processed: one batch was digested by in-gel digestion, while the second batch was split between in-gel and on-bead digestion.

In-gel digestion: Proteins were eluted with 0.1 M glycine (pH 2.2) and neutralized with Tris-HCl (pH 8.0). After gel separation, each gel band was reduced with 10 mM DTT for 30 minutes at 60 °C and alkylated with 20 mM iodoacetamide for 45 minutes at room temperature in the dark. Proteins were digested overnight at 37 °C with sequencing-grade trypsin (Thermo Scientific, Cat# 90058). Peptides were extracted twice with 5% formic acid and 60% acetonitrile, then dried under vacuum.

On-bead digestion: Beads were washed multiple times with 50 mM NH₄HCO₃ until detergent levels were reduced to <0.4%. Washed beads were incubated with 0.2 µg of trypsin in 20 µL of 50 mM NH₄HCO₃ and 0.01% Protease Max (Promega) at 37 °C for 2 hours. An additional 0.2 µg of trypsin was added and incubated for another 2 hours at 37 °C. The solution was separated from the beads, acidified to pH 3 with 10% formic acid, and desalted using a stage tip before LC-MS/MS analysis. HCRG1 was not detected after on-bead digestion, likely due to direct binding to the antibody.

#### LC-MS/MS analysis

Samples were analyzed using a Nano LC-MS/MS system (Dionex Ultimate 3000 RLSCnano System, ThermoFisher) coupled with an Orbitrap Eclipse mass spectrometer (ThermoFisher). Three microliters of digested sample were loaded onto a fused silica trap column (Acclaim PepMap 100, 75 µm × 2 cm, ThermoFisher) and washed for 5 minutes at 5 µL/min with 0.1% TFA. The trap column was connected to an analytical column (NanoEase MZ peptide BEH C18, 130 Å, 1.7 µm, 75 µm × 250 mm, Waters) for peptide separation at 300 nL/min using a segmented linear gradient (4–15% B in 30 min, 15–25% B in 40 min, 25–50% B in 44 min, and 50–90% B in 11 min).

MS1 spectra were acquired using Orbitrap analysis at a resolution of 120,000 with a scan range from m/z 375–1500, an AGC target of 1E6, and a maximum injection time of 100 ms. A top-3-second duty cycle scheme was used, selecting parent ions with charge states of 2–7 and applying dynamic exclusion for 60 seconds. Parent ions were isolated in the quadrupole with a 1.2 m/z window and fragmented by higher-energy collisional dissociation (30% normalized collision energy). Fragments were analyzed in the Orbitrap at a resolution of 15,000.

#### Data processing

LC-MS/MS peak lists were generated using Thermo Proteome Discoverer (v2.4) into MASCOT Generic Format (MGF) and searched against the H. saltator protein database (NCBI_GCF_003227715.2_Hsal_v8.6_protein.fasta) and a contaminant database (CRAP) using X!Tandem (GPM Fury). Search parameters included a parent mass tolerance of ±7 ppm, a fragment mass tolerance of ±20 ppm, variable modifications (methionine oxidation, asparagine deamination, tryptophan oxidation, dioxidation, methionine dioxidation, glutamine to pyro-glutamine), and trypsin cleavage specificity (up to one missed cleavage). The peptide false discovery rate (FDR) was 0.07%. Results were quantified using the R package IPinquiry4.

### Ortholog identification and codon-based evolutionary analysis

Orthologous groups were inferred using OrthoFinder v2.5.5 from a curated 70-species protein dataset with a supplied rooted species tree. For NCBI-derived proteomes, the longest isoform per gene was retained; GAGA representative proteomes and FigShare one-transcript-per-gene annotations were used as provided. The 70-species working tree was generated by grafting the dated ant phylogeny from Vizueta et al. (2025) onto a TimeTree-derived insect backbone. The ant subtree was inserted at the most recent common ancestor of ant taxa in the backbone and rescaled to match the corresponding ant crown age.

Co-orthologs of HCRG1 (XP_011135446.1) and its in-paralog (XP_011135445.1), as well as co-orthologs of Hex70c (XP_011142758.1), were extracted from the OrthoFinder results. CDS sequences were retrieved from species-specific CDS files using OrthoFinder protein identifiers. Sequences with frame disruption or internal stop codons were removed, and protein entries inconsistent with their CDS translations were replaced by CDS-derived translations. Protein sequences were aligned using MAFFT v7.525, and codon alignments were generated using PAL2NAL v14. Codon alignments were converted to sequential PHYLIP format, and OrthoFinder gene trees were pruned to match the retained sequences.

Codon-based evolutionary analyses were performed using codeml in PAML v4.10.10. Site models M0, M1a, M2a, M7, and M8 were run for each gene. Branch models were performed using two-ratio and multi-ratio settings to test lineage-, taxon-, and trait-associated variation in dN/dS. Two-ratio branch models used model = 2 and NSsites = 0, with foreground branches labeled in the gene tree. Branch-site models used model = 2 and NSsites = 2, comparing the alternative model (fix_omega = 0) against the null model with ω fixed at 1 (fix_omega = 1, omega = 1). Likelihood-ratio tests were calculated as 2ΔlnL and assessed using chi-square tests.

### AlphaFold3 structural modeling and ligand docking analysis

For structural prediction, AlphaFold3 (AF3, version 3.0.1) was used to model four protein systems: the *Harpegnathos saltator* Hex70c hexamer, the HCRG1-bound Hex70c hexamer, the *Drosophila melanogaster* Lsp2 hexamer, and the HCRG1-bound DmLsp2 hexamer. For the *H. saltator* models, six copies of the inferred secreted form of Hex70c (XP_011142758.1, residues 36–682) were used to model the Hex70c hexamer, either alone or together with six copies of the inferred secreted form of HCRG1 (XP_011135446.1, residues 19–82) ^83^. For the DmLsp2 models, six copies of the inferred mature form of DmLsp2/Lsp2 isoform B (NP_001287042.1, residues 22–701) were used to model the DmLsp2 hexamer, either alone or together with six copies of the inferred secreted form of HCRG1. Each protein copy was defined as an independent chain in the AF3 input. For each target, an initial no-template AF3 search was performed using full MSA features and 20 independent random seeds. The resulting models were ranked according to AF3 confidence scores, primarily the model ranking score, predicted TM-score (pTM), and inter-chain predicted TM-score (ipTM). To further optimize the hexameric core conformations used for downstream docking, we performed an iterative rolling-template refinement strategy. In this procedure, the best model from the Hex70c or DmLsp2 hexamer was used as a chain-specific custom template to guide modeling of the corresponding HCRG1-bound complex. The best HCRG1-bound complex model was then used to extract the Hex70c or DmLsp2 hexameric core as templates for the next hexamer-only round. This hexamer/HCRG1-bound complex-guided refinement cycle was repeated until no further improvement in ranking score, pTM, or ipTM was observed. The best-scoring models were selected for downstream docking analysis.

Ligand structures were prepared for three juvenile hormone-related compounds: juvenile hormone III (JH III; PubChem CID: 5281523), juvenile hormone bisepoxide (JHB3; PubChem CID: 6439054), and methyl farnesoate (MF; PubChem CID: 5275508). Three-dimensional ligand structures were downloaded in SDF format, and both ligand SDF files and AF3-derived protein receptor mmCIF files were converted for docking using Open Babel 3.1.0. Protein receptor structures were further cleaned to retain protein ATOM records. Receptor and ligand PDBQT files were prepared using MGLTools 1.5.7, including the addition of hydrogens and assignment of Gasteiger charges.

Molecular docking was performed using AutoDock Vina (v1.2.6). As a benchmark, the experimentally determined JH III-binding pose from the solution NMR structure PDB ID: 2RQF ^28^ was simulated using Vina. For AlphaFold3-derived models, grid boxes were defined and iteratively refined using a global-to-local best-pose docking strategy. The final docking was performed using a uniform 24 Å cubic grid box centered on the best observed pose for each receptor–ligand combination, with exhaustiveness set to 512 and 20 independent runs per condition using randomized seeds. The top-scoring model from each run was retained for downstream plotting and comparative analysis.

#### Structural metrics assessment

Ligand RMSD was calculated after chain-permutation-aware Cα alignment of Hex70c backbones. JH III heavy-atom RMSD was calculated using the Hex70c-alone mapped-site pose as the reference.

Binding pocket volume was calculated using PyVOL v1.7.8 with MSMS v2.6.1. PyVOL was run on a local pocket region containing protein residues within 12 Å of the docked JH III pose. The ligand centroid was used as the pocket-defining coordinate. A solvent-excluded surface model with a 1.4 Å probe radius was used.

Contacting residues were defined as receptor residues with any heavy atom within 4.0 Å of any JH III heavy atom.

Buried surface area was calculated using FreeSASA v2.2.1 as SASA(protein) + SASA(JH III) − SASA(complex). Ligand carbon and oxygen atoms were assigned standard element-based radii.

Putative hydrogen bonds were defined using a 3.6 Å heavy-atom distance cutoff between JH III heteroatoms and receptor N/O/S atoms. No angular cutoff was applied.

Hydrophobic contacts were defined as Ala, Val, Leu, Ile, Met, Phe, Trp, Pro, and Tyr residues with any heavy atom within 4.0 Å of JH III.

### Bulk RNA sequencing

Two intact non-duelers, two intact duelers, two GFP RNAi duelers, and two HCRG1 RNAi duelers from a single experimental colony were collected for bulk RNA sequencing. For each individual, the brain, abdominal fat body, and ovary were dissected, followed by RNA extraction and library preparation. RNA isolation, including tissue dissection and chloroform extraction, followed the same protocol as for qPCR sample preparation. The aqueous phase was carefully collected and purified using RNA Clean & Concentrator Kits (Zymo Research, Cat# RRC-5) following the manufacturer’s instructions. RNA quality and concentration were assessed using a Qubit 4 Fluorometer (Invitrogen).

High-quality RNA was used for library preparation with the NEBNext® Ultra II Directional RNA Library Prep Kit according to the manufacturer’s guidelines. RNA sequencing reads were aligned to the *Harpegnathos* genome (version 8.5) using STAR (v2.7.11). Read alignment data were processed with featureCounts to generate count files, which were analyzed using the R package DESeq2 for differential gene expression (DEG) analysis. DESeq2-normalized counts were used for downstream analyses, including transcriptomic distance (TSD) measurements and Gene Set Enrichment Analysis (GSEA). TSD was calculated using both Jensen-Shannon Divergence and Rankings Correlation Distance (Manatakis et al., 2021). GSEA was performed using the R package *clusterProfiler* with a gene set file created using GMT Helper (https://biit.cs.ut.ee/gmt-helper/).

### Fly survival assay

*UAS-HCRG1* transgenic flies were generated by cloning *Harpegnathos HCRG1* cDNA into the pUAST-attB vector, followed by microinjection into fly embryos by BestGene. *UAS-EGFP* (BL#5428) and *UAS-dFOXO* (BL#42221) were obtained from the Bloomington *Drosophila* Stock Center. To achieve precise spatiotemporal control, all Gal4 drivers were combined with tub-Gal80^ts^. Virgin female Gal4 flies were crossed with male UAS-HCRG1 flies and raised at 18 °C until F1 progeny emerged. Virgin F1 females with the desired genotype were collected and transferred in groups of approximately 30 flies per vial. F1 flies were maintained at 29 °C, and survival was monitored daily until all individuals had died. Food vials were replaced three times per week to prevent premature death caused by food spoilage or poor vial conditions.

### Multivariate correlation analysis

Pearson correlation analyses were performed using the Hmisc R package. Correlation coefficients and corresponding *p*-values were visualized using corrplot, with significance levels annotated. Flattened correlation matrices for figure annotation were generated using base R.

### Quantification and statistical analysis

Statistical analyses, including unpaired *t*-test and one-way ANOVA with Tukey’s multiple comparisons test, were performed using R scripts. The value of n, mean ± SEM, and *p*-values are reported in the Results, Figures, and Figure Legends. All samples used for parametric tests passed the Normality test.

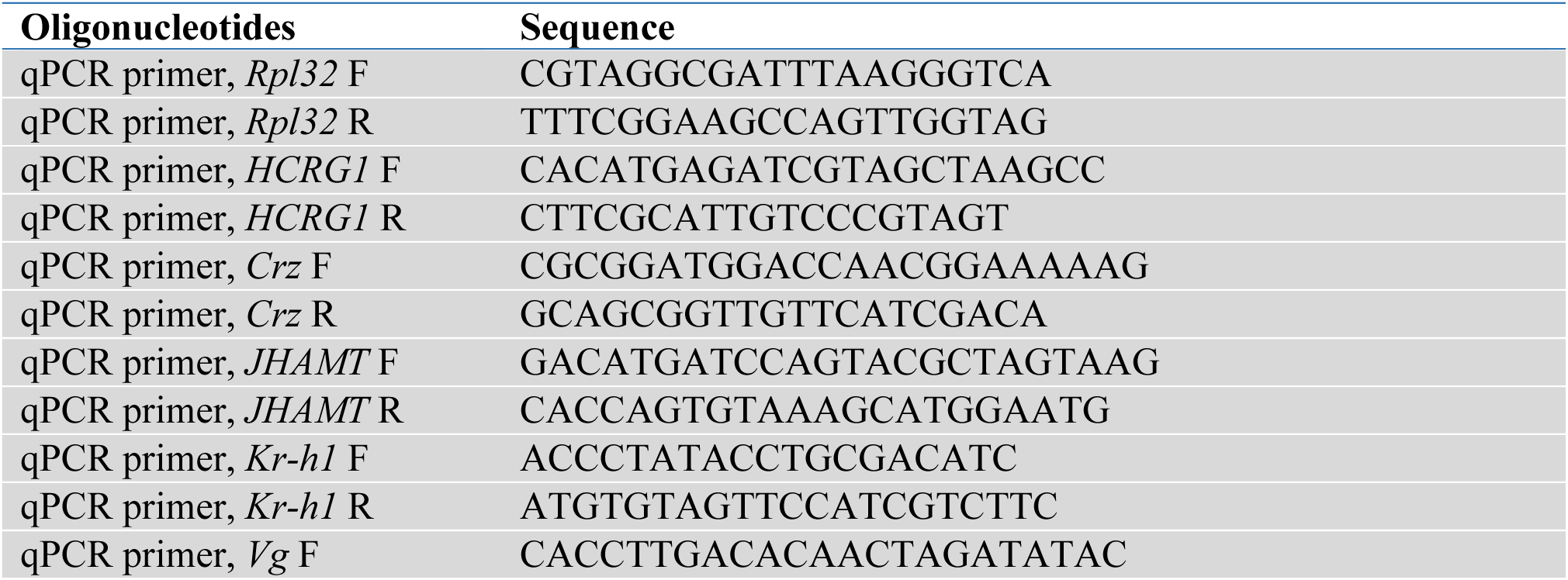
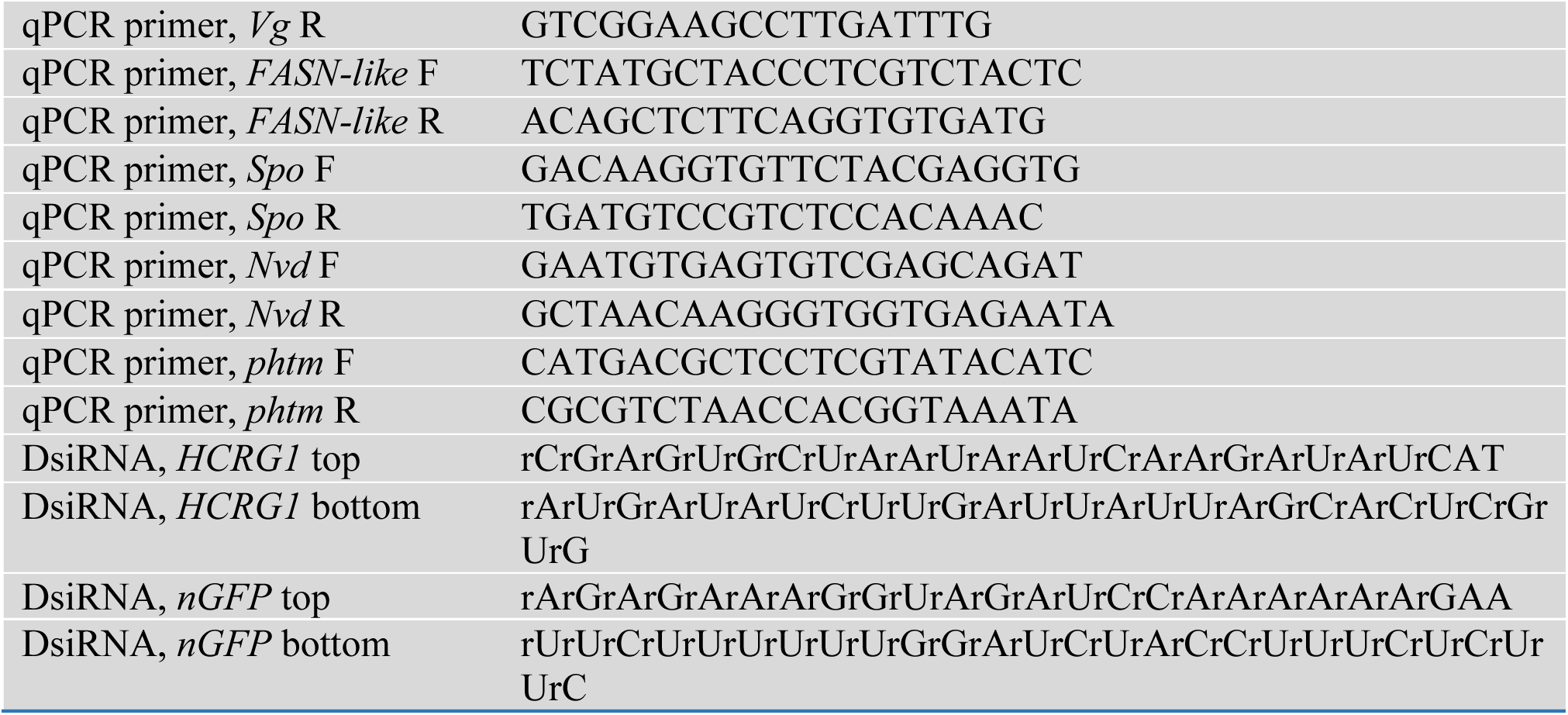

---

**Extended Data Fig. 1.**
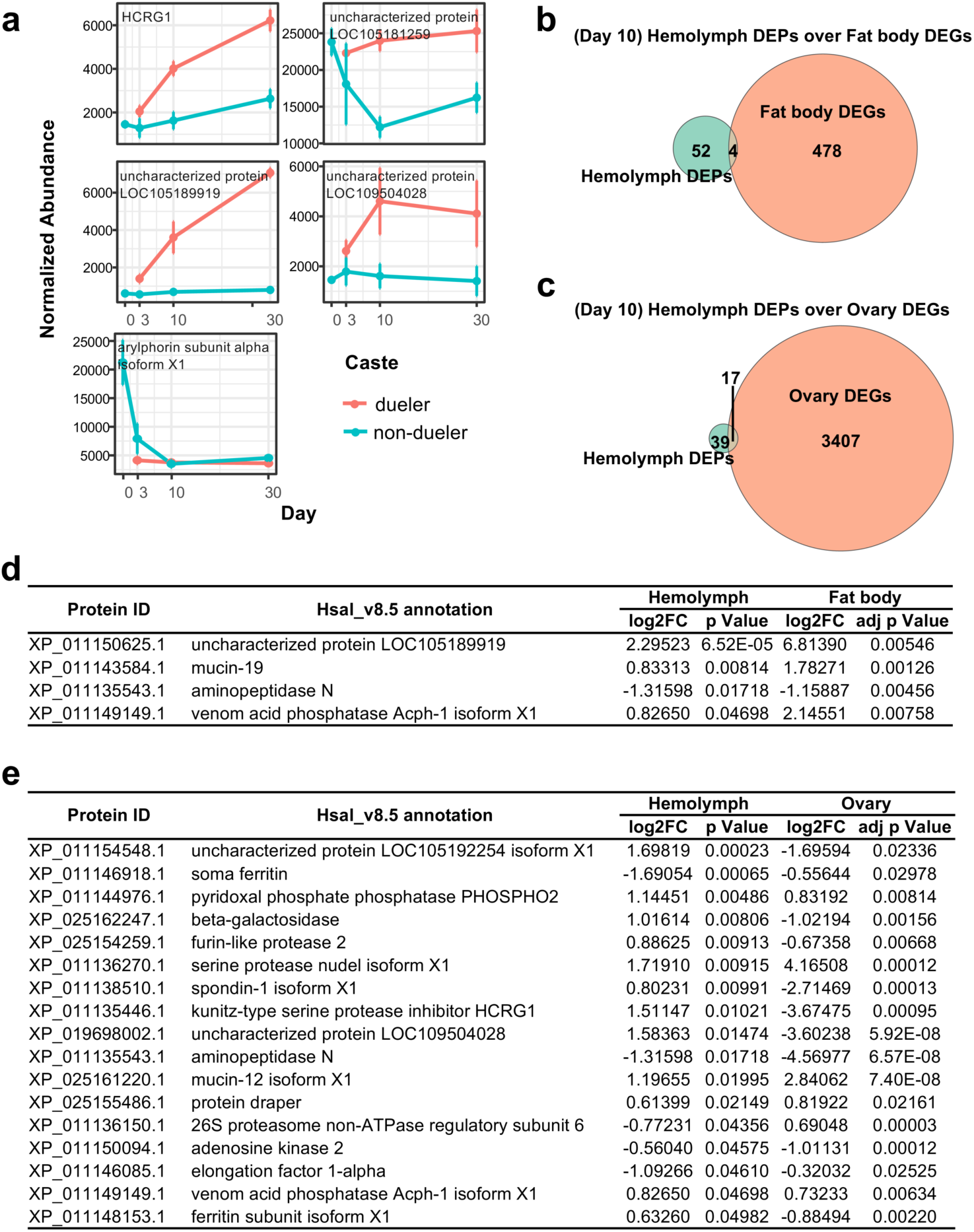
Shared factors between transcriptomes and hemolymph proteome. **a,** Curve plots of the four shared proteins from Fig. 1b and arylphorin subunit alpha isoform X1, showing their circulating levels throughout the transition. Data are from proteomic analysis. Error bars represent SEM. **b,** Venn diagram comparing the abdominal fat body transcriptome and hemolymph proteome at day 10. **c,** Venn diagram comparing the ovary transcriptome and hemolymph proteome at day 10. **d,** List of the four shared factors from **b**. **e,** List of the 17 shared factors from **c**.

**Extended Data Fig. 2.**
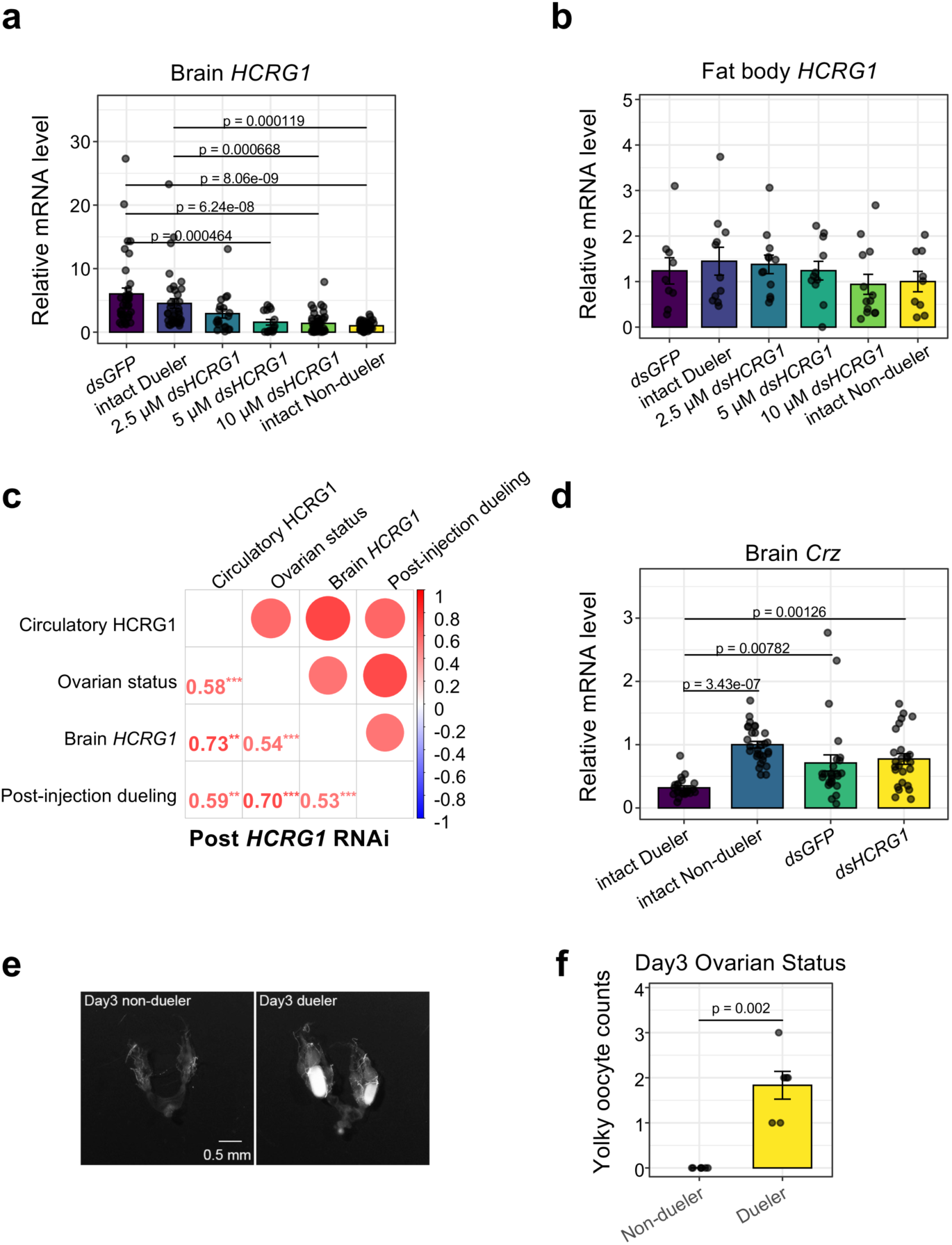
Effects of HCRG1 KD on gene expression and phenotypic markers. **a,b,** qPCR analysis of HCRG1 expression in brains (**a**) and fat bodies (**b**) from different treatment groups. RPL32 was used as an internal control. Results from multiple colonies were combined, and each expression value was normalized to the average of non-duelers from the same colony. Error bars indicate SEM. Statistical significance was determined by one-way ANOVA followed by Tukey’s post hoc analysis. Sample sizes: **a,** dsGFP (n = 37), intact duelers (n = 37), 2.5 μM dsHCRG1 (n = 19), 5 μM dsHCRG1 (n = 15), 10 μM dsHCRG1 (n = 47), and intact non-duelers (n = 43); **b,** dsGFP (n = 9), intact duelers (n = 11), 2.5 μM dsHCRG1 (n = 12), 5 μM dsHCRG1 (n = 11), 10 μM dsHCRG1 (n = 13), and intact non-duelers (n = 9). **c,** Correlation analysis of hemolymph HCRG1 protein levels, ovarian development, brain HCRG1 mRNA levels, and post-treatment dueling activity in ants from HCRG1 RNAi experimental colonies. Circle size represents the magnitude of the absolute correlation coefficient. Numbers denote correlation coefficient values. Color scale: blue indicates negative correlation; red indicates positive correlation. **d,** qPCR analysis of Crz expression in brains from different treatment groups. RPL32 was used as an internal control. Results from multiple colonies were combined, and each expression value was normalized to the average of non-duelers from the same colony. Error bars represent SEM. p-values were calculated using one-way ANOVA with Tukey’s post-hoc analysis. Sample sizes: intact duelers (n = 23), intact non-duelers (n = 29), dsGFP (n = 25), dsHCRG1 (10 μM, n = 26). **e,** Representative images of dissected ovaries of day 3 non-duelers and duelers (scale bar = 0.5 mm). **f,** Quantification of yolky oocytes. Error bars represent SEM. p-values were calculated using t-test.

**Extended Data Fig. 3.**
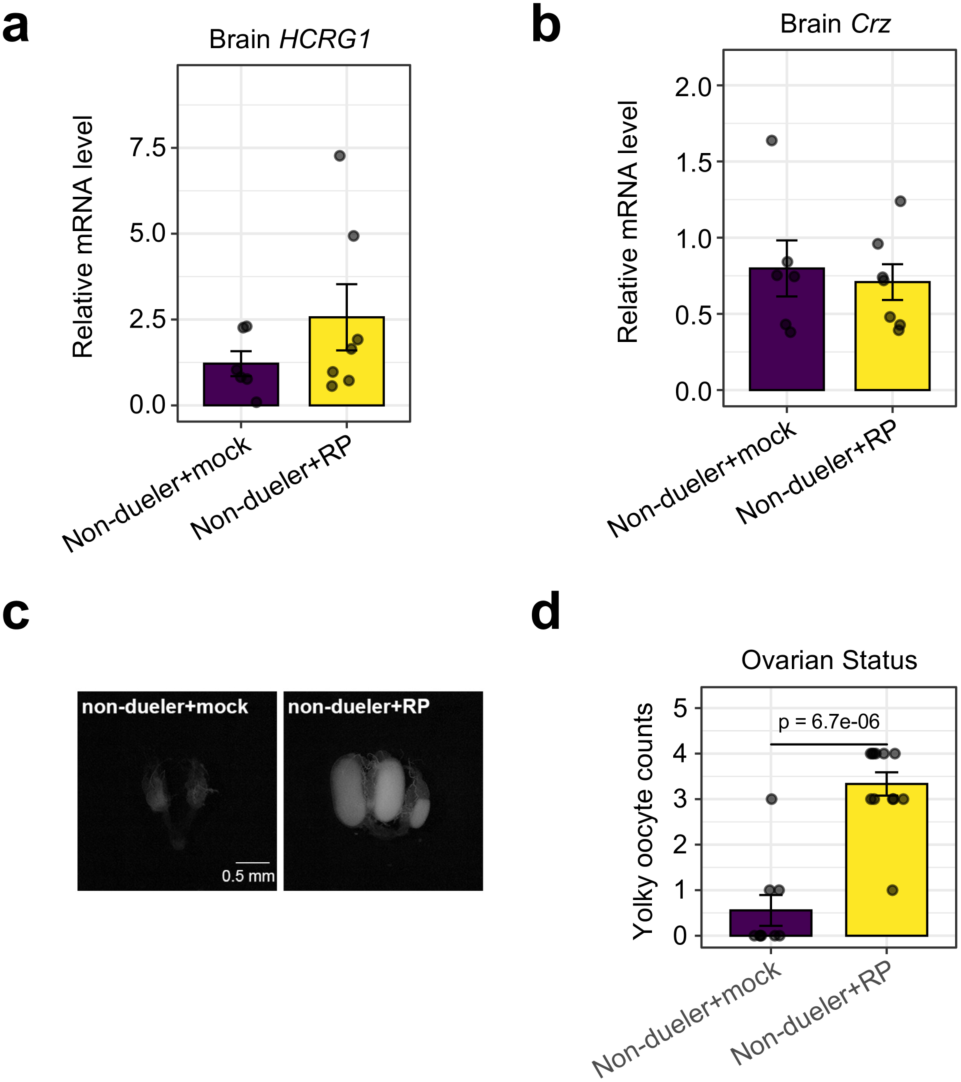
Ectopic HCRG1 induces partial gamergate-traits formation in non-duelers. **a,b,** qPCR analysis of HCRG1 (**a**) and Crz (**b**) expression in brains of non-duelers injected abdominally with HBSS (mock, n=6) or RP (n=7). RPL32 was used as an internal control. Error bars represent SEM. **c,** Representative images of dissected ovaries from non-duelers injected with abdominal HBSS (mock, n=9) or RP (n=12). Scale bar = 0.5 mm. **d,** Quantification of yolky oocytes. Error bars represent SEM. p-values were calculated using t-test.

**Extended Data Fig. 4.**
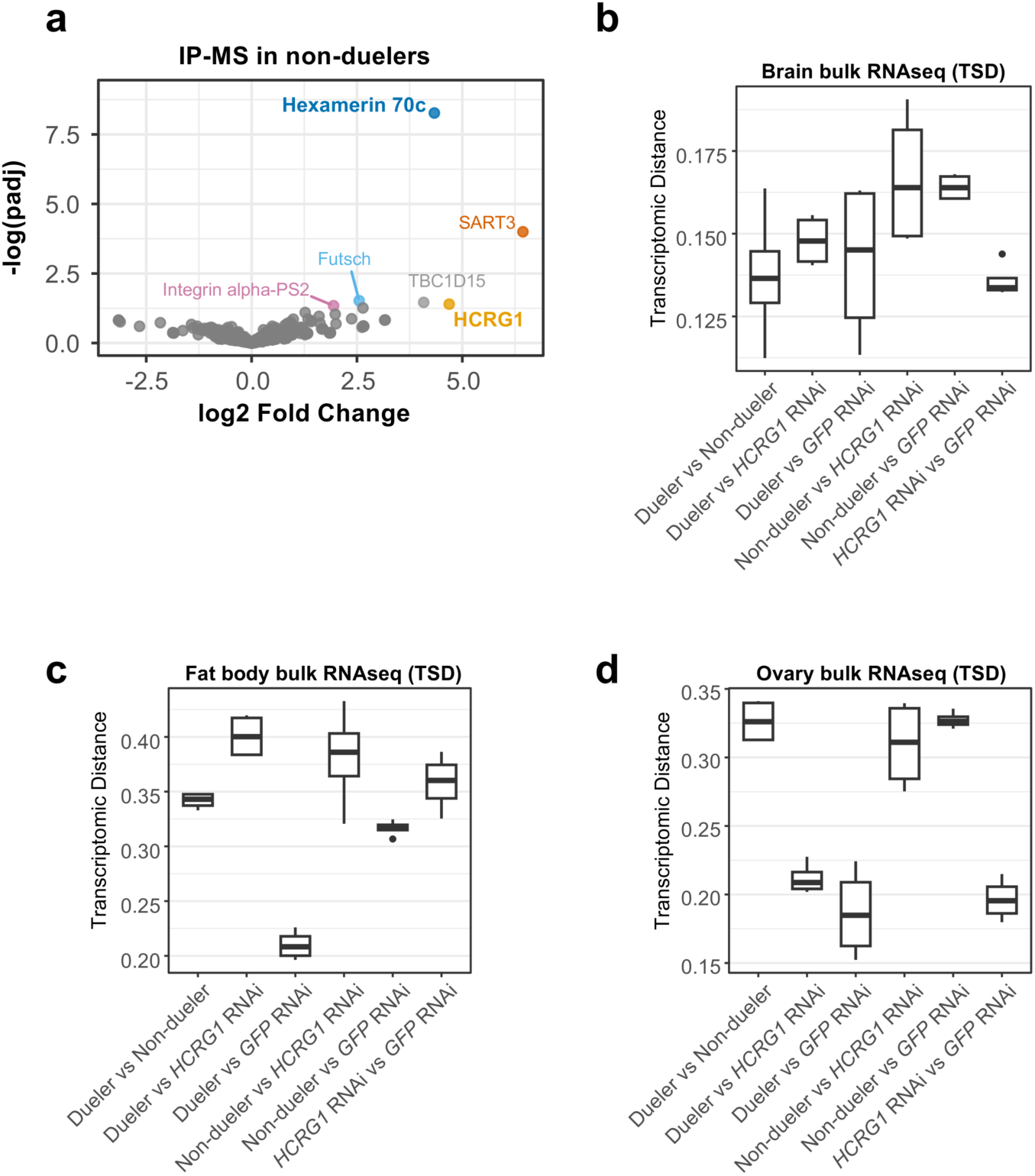
Analysis of proteins interacting with HCRG1 and transcriptomic distance across different treatments. **a,** Volcano plot of IP-MS results from non-duelers. Proteins enriched in HCRG1 antibody-crosslinked beads were compared to the IgG control. Each dot represents an identified protein. Proteins significantly enriched in the HCRG1 pulldown (adjusted p < 0.05) are highlighted in color: Hexamerin 70c (blue), HCRG1 (orange), SART3 (vermilion), Futsch (light blue), TBC1D15 (gray), and Integrin alpha-PS2 (magenta). Non-significant proteins (p > 0.05) are shown in gray. **b-d,** Box plots of transcriptomic distance (TSD) calculated from bulk RNA-seq data in the brain (**b**), fat body (**c**), and ovary (**d**), comparing the following groups: intact duelers, intact non-duelers, GFP RNAi controls, and HCRG1 RNAi-treated individuals.

**Extended Data Fig. 5.**
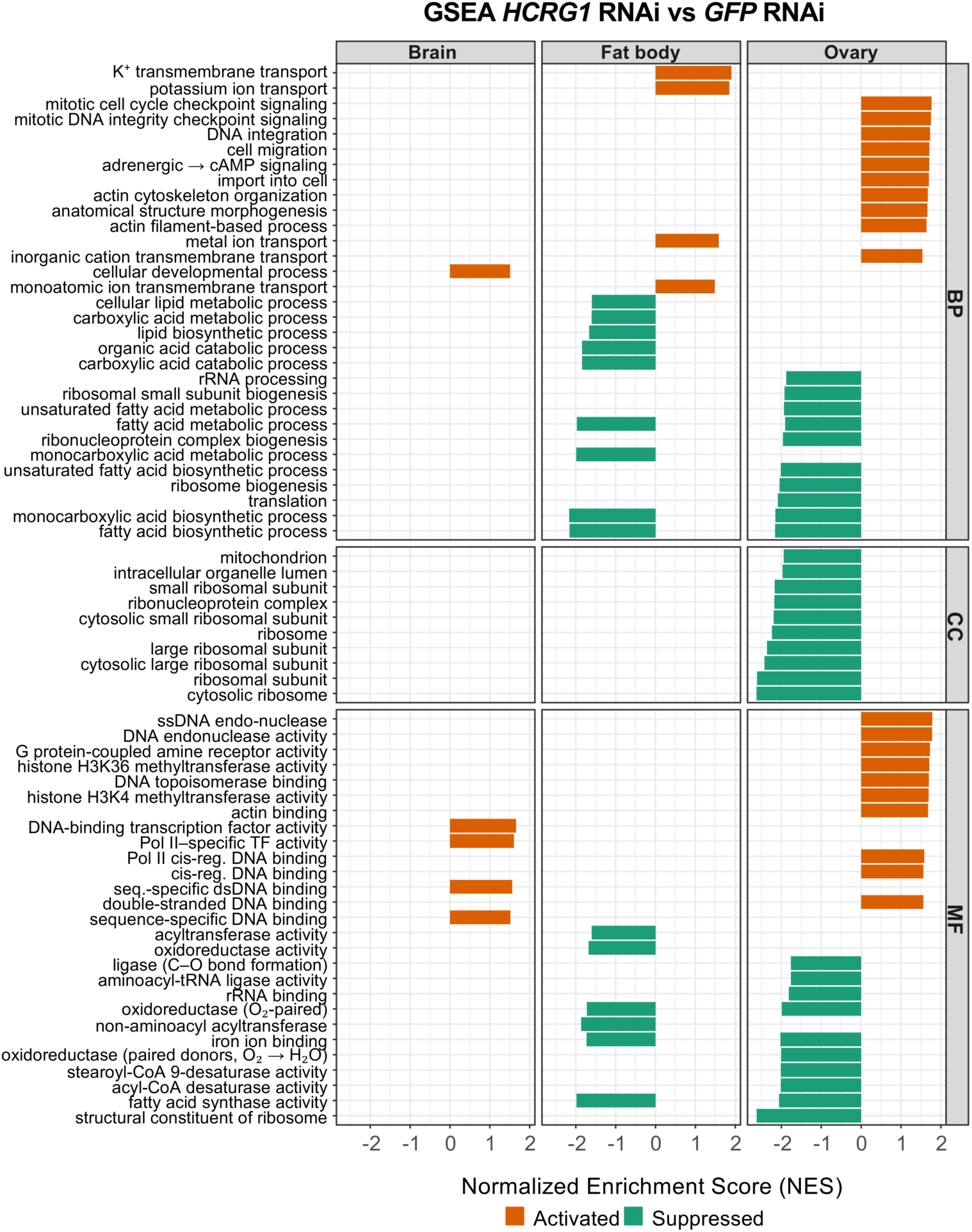
GSEA of DEGs in brain, fat body, and ovary from *HCRG1* RNAi vs. *GFP* RNAi. GSEA of differentially expressed genes (DEGs) comparing HCRG1 RNAi to GFP RNAi in brain, fat body, and ovary tissues. Each bar represents a significantly enriched Gene Ontology (GO) term (FDR < 0.05), with the horizontal axis indicating the normalized enrichment score (NES). Bars extending to the right (orange) indicate positively enriched gene sets (activated), while bars extending to the left (green) indicate negatively enriched gene sets (suppressed). GO terms are grouped by ontology: BP (Biological Process), CC (Cellular Component), and MF (Molecular Function). Long GO term names have been abbreviated for clarity.

**Extended Data Fig. 6.**
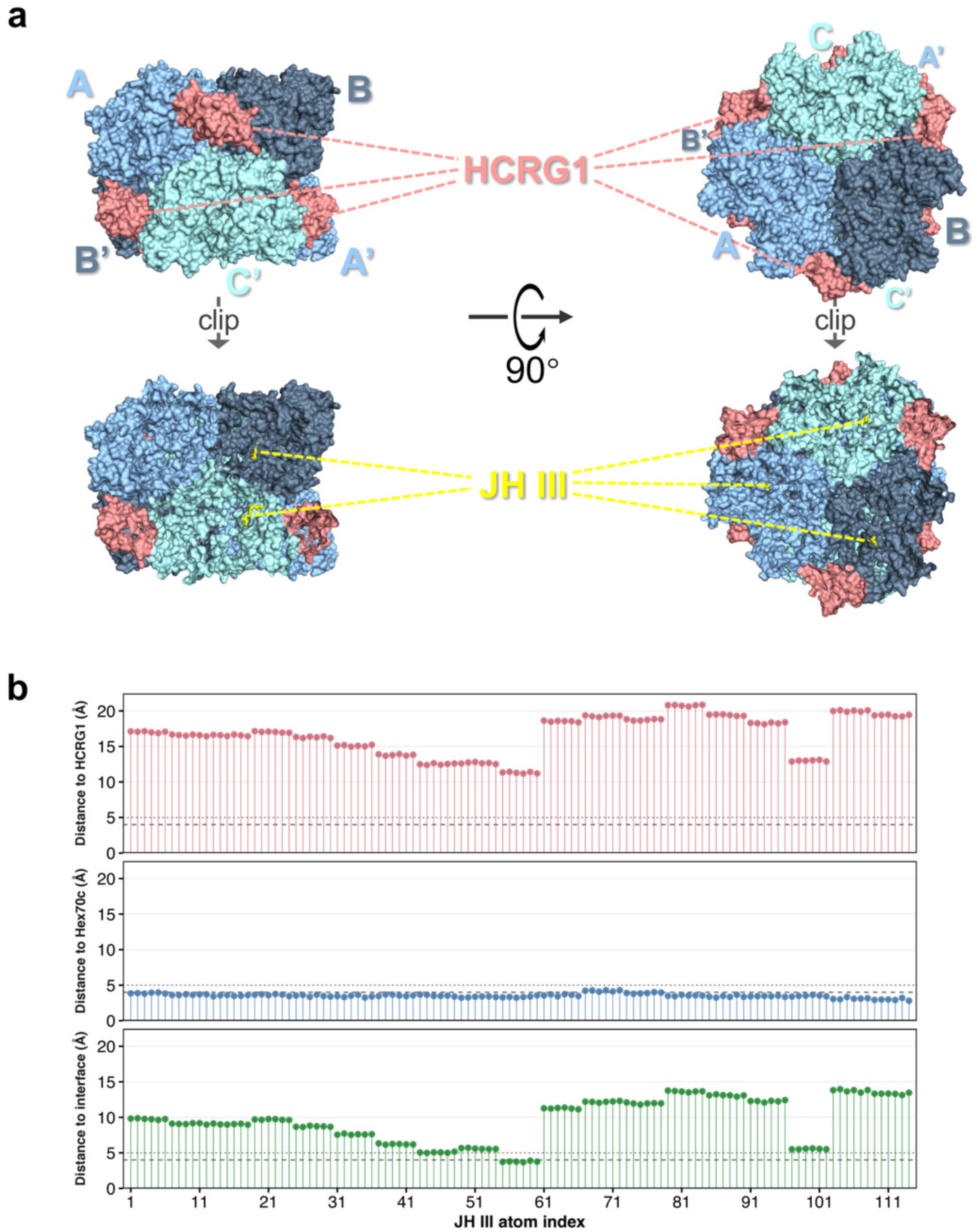
HCRG1-bound Hex70c forms internal JH III-binding pockets. **a,** Structural organization and internal ligand positioning of the HCRG1–Hex70c complex. The upper panels show surface-rendered models of the predicted HCRG1-Hex70c complex (AF3 model) in side view (left) and top view (right). The hexameric Hex70c core consists of six subunits arranged in two staggered layers, each comprising three radially distributed chains. The upper layer is labeled A (pale cyan), B (slate), and C (light blue), with the corresponding subunits in the lower layer labeled A′, B′, and C′. Six HCRG1 subunits (salmon) partially surround the periphery of the complex. The lower panels display clipped views that expose the internal ligand-binding cavities, each occupied by a JH III molecule (yellow sticks) docked using AutoDock Vina. Internal cartoon representations are overlaid within each subunit and colored to match the surface. **b,** Nearest-distance analysis of six symmetry-mapped docked JH III molecules in the HCRG1-bound Hex70c hexamer. Each lollipop represents one JH III heavy atom and shows its nearest heavy-atom distance to HCRG1, Hex70c, or the HCRG1–Hex70c interface. Interface residues were defined as residues with at least one heavy atom within 5.0 Å of the opposing partner. Dashed and dotted lines mark the 4 Å and 5 Å thresholds. No JH III atom was within 5 Å of HCRG1, whereas all 114 atoms were within 5 Å of Hex70c. Atoms close to the interface mainly reflect proximity to Hex70c-side interface residues rather than direct contact with HCRG1.

**Extended Data Fig. 7.**
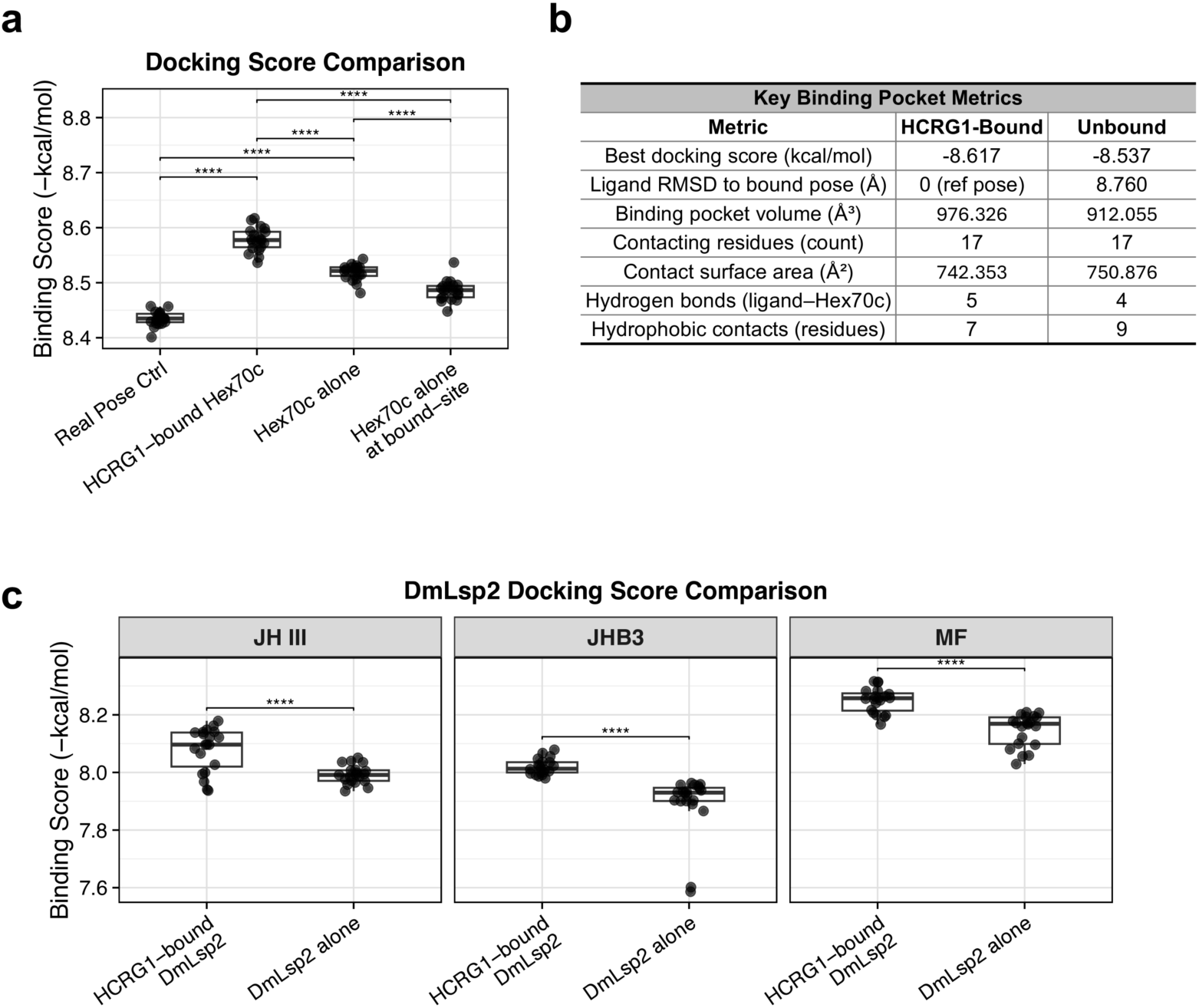
Docking-based comparison of JH-related ligand binding to Hexamerin complexes. **a,** Boxplot of docking score comparison of JH III across different structural contexts. AutoDock Vina scores are plotted as transformed binding scores, calculated as −1 × Vina score, so higher values indicate stronger predicted binding. The real pose control represents redocking of JH III into the native pocket of the 2RQF juvenile hormone-binding protein structure. HCRG1-bound Hex70c and Hex70c alone represent focused JH III docking to the best pose centers identified in the corresponding receptor models. “Hex70c alone at bound-site” represents docking to Hex70c alone using the best JH III docking site from the HCRG1-bound Hex70c model after structural mapping onto the Hex70c-alone receptor. Each point represents one independent docking seed. Boxes indicate the interquartile range with the median line. Statistical significance was assessed by one-way ANOVA followed by Tukey’s HSD post hoc test. ****P ≤ 0.0001. **b,** Structural metrics comparing JH III binding to Hex70c in the presence and absence of HCRG1. The ligand pose was compared across two structural conditions (HCRG1-bound vs. unbound Hex70c), and multiple interaction features were quantified. Ligand RMSD was measured after aligning Hex70c backbones. **c,** Ligand-specific docking score comparison between HCRG1-bound DmLsp2 and DmLsp2 alone for JH III, JHB3, and methyl farnesoate (MF). Scores are plotted as transformed binding scores, calculated as −1 × Vina score. Each point represents one independent docking seed. Boxes indicate the interquartile range with the median line. Statistical significance was assessed using an unpaired two-sided t-test for each ligand comparison. ****P ≤ 0.0001.

**Extended Data Fig. 8.**
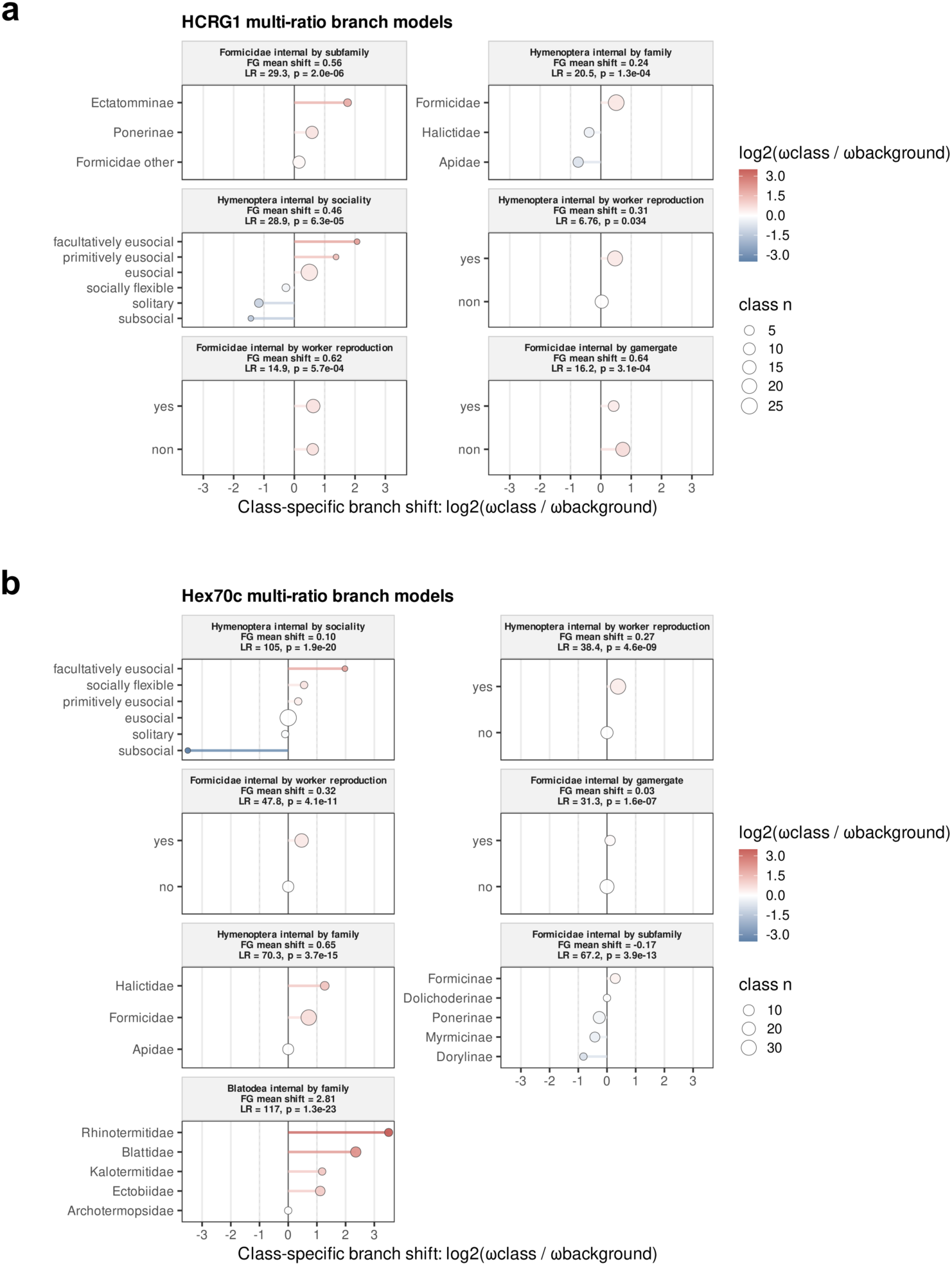
Multi-ratio branch models for HCRG1 and Hex70c. **a,b,** Centered lollipop plots show class-specific ω shifts from multi-ratio branch models for HCRG1 (a) and Hex70c (b). Each facet represents one model in which a broader foreground lineage was subdivided by taxonomic or social-trait class. The x axis shows log₂(ω_class/ω_background), with negative values indicating reduced class-specific ω and positive values indicating elevated class-specific ω relative to the model-specific background. Color represents the magnitude and direction of this class-level shift, and point size indicates class size. Facet headers indicate the tested foreground subdivision, the weighted mean shift across non-background classes, and the model-level LR and p value for the multi-ratio model versus the corresponding one-ratio model.

**Extended Data Fig. 9.**
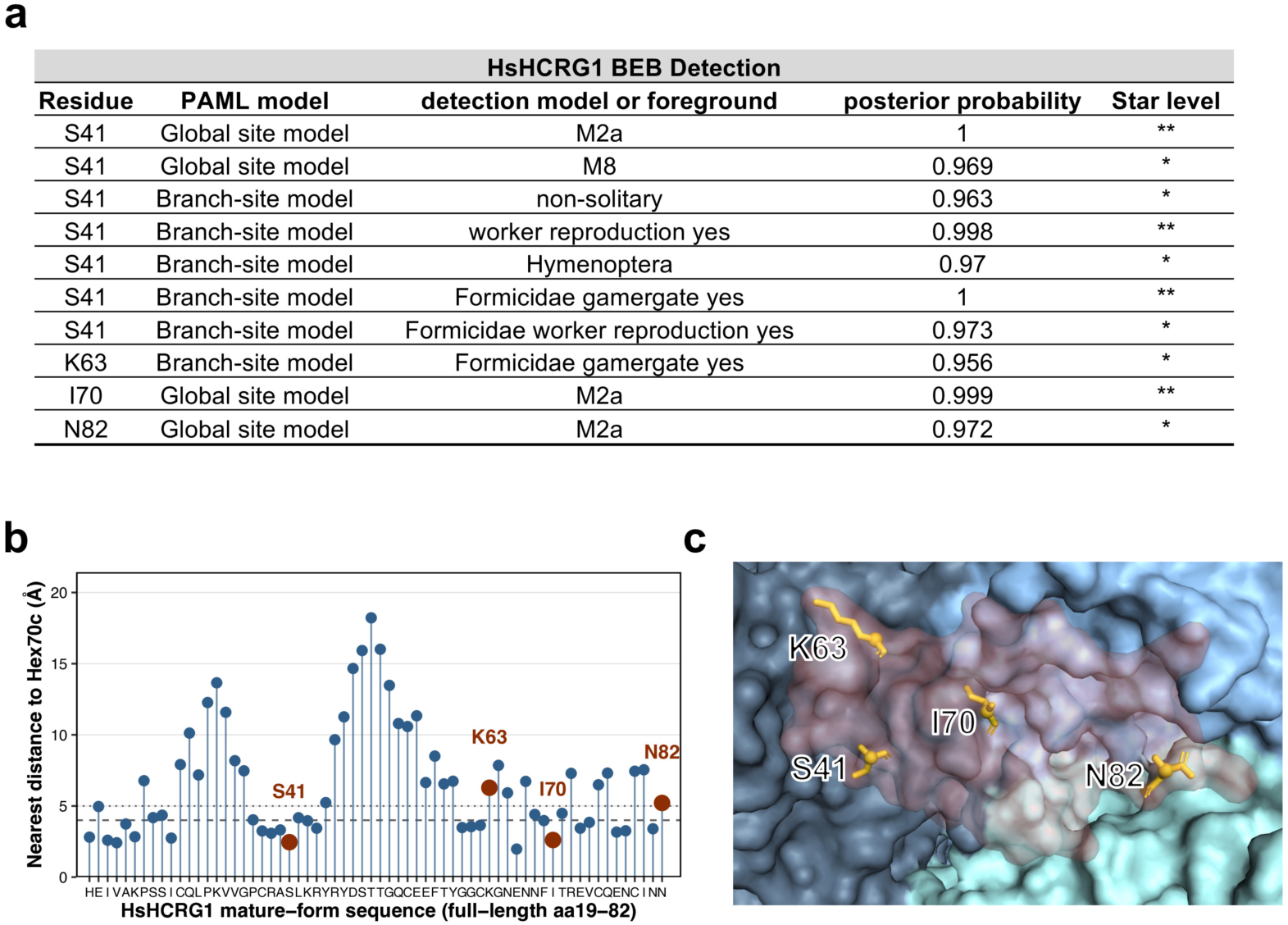
Evolutionary support and structural localization of reportable HCRG1 BEB sites. **a,** Summary of reportable Bayes empirical Bayes (BEB) sites detected in HsHCRG1. The table lists the HCRG1 residue, PAML model class, detection model or branch-site foreground, posterior probability, and BEB support level. Only BEB sites with posterior probability ≥ 0.95 and mappable to non-gap HsHCRG1 residues are shown. Global site-model support includes M2a and M8 results, whereas branch-site evidence is shown for foreground tests containing HsHCRG1. Single and double asterisks indicate posterior probabilities ≥ 0.95 and ≥ 0.99, respectively. S41 showed the strongest cross-model support, being detected by both global site models and multiple Hs-containing branch-site foregrounds, whereas K63, I70, and N82 were supported by individual model classes or foregrounds. **b,** Lollipop plot showing the nearest heavy-atom distance from each mature-form HCRG1 residue to Hex70c in the best HCRG1-bound Hex70c structural model. The x-axis shows the mature HCRG1 peptide sequence corresponding to full-length HsHCRG1 aa19–82, and the y-axis shows the minimum distance to any Hex70c heavy atom. For each sequence position, the plotted value represents the minimum distance across the six HCRG1 copies in the complex. Reportable BEB sites are highlighted in brown and labeled. Dashed and dotted horizontal lines indicate 4 Å and 5 Å distance thresholds, respectively. **c,** Structural view of the reportable HCRG1 BEB sites in the HCRG1-bound Hex70c complex. The six Hex70c subunits are shown as solid surfaces in three alternating colors: light blue, slate blue, and cyan. HCRG1 chains are shown as semi-transparent salmon surfaces to reveal the underlying Hex70c interface. Reportable BEB residues S41, K63, I70, and N82 are highlighted as yellow sticks and labeled. Residue numbering corresponds to full-length HsHCRG1, whereas the modeled mature HCRG1 peptide spans aa19–82. The highlighted sites occupy distinct positions across the HCRG1 surface, with S41 and I70 positioned closer to Hex70c-contacting regions and K63 and N82 located more peripherally.

**Extended Data Fig. 10.**
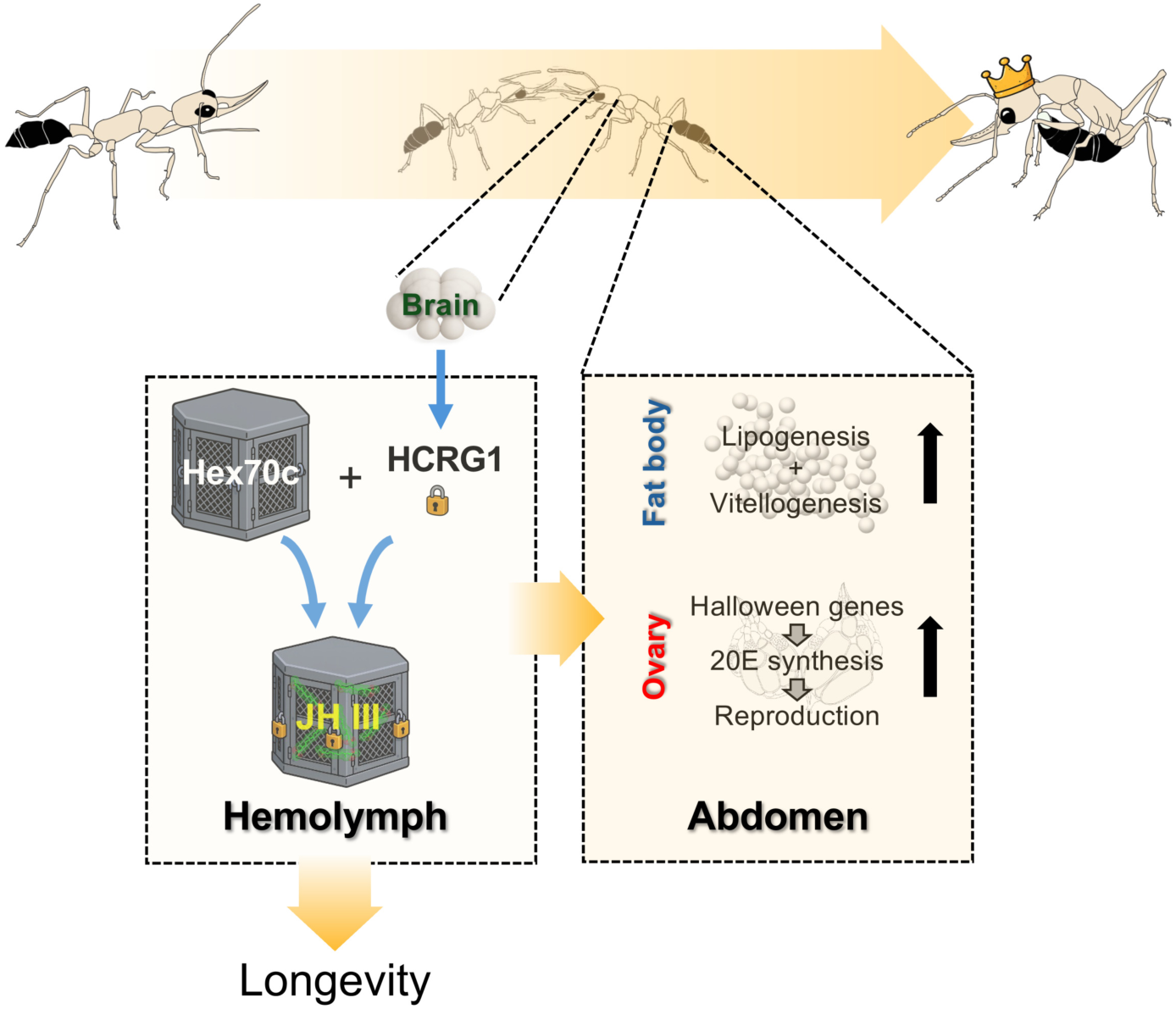
Graphical Summary: HCRG1-mediated JH sequestration promoting caste in Harpegnathos ants. In response to queen loss, Harpegnathos workers engage in antennal dueling, initiating their transition into reproductive pseudo-queens (gamergates). During this process, the brain secretes HCRG1, which binds to the Hex70c in the hemolymph. This interaction locks the cage-like Hex70c structure, enhancing its capacity to sequester JH III and thereby reducing free JH levels in circulation. The resulting JH depletion triggers systemic physiological reprogramming in the abdomen, including increased lipogenesis and vitellogenesis in the fat body, as well as activation of Halloween gene-dependent 20E synthesis and reproductive maturation in the ovary. In parallel, suppression of JH signaling promotes lifespan extension. Together, these endocrine changes support the formation of the reproductive and long-lived gamergate phenotype.

